# GeneBench: Assessing AI Agents for Multi-Stage Inference Problems in Genomics and Quantitative Biology

**DOI:** 10.64898/2026.04.22.720113

**Authors:** Jeremy Li, Andrew Ho

## Abstract

We introduce GeneBench, a benchmark for AI agents on realistic multi-stage scientific data analysis in genetics and quantitative biology. Existing biology benchmarks mostly measure knowledge retrieval, execution of routine pipelines, or a single analysis step. Yet they do not capture the broader scope of work that occupies much of computational scientists’ time: cleaning and normalizing assay, phenotype, or clinical data; exploratory data analysis; statistical model selection and diagnostic iteration; and producing a conclusion that informs a downstream scientific or translational decision. GeneBench addresses this gap with 103 evaluations targeting quantities of direct practical relevance across 10 domains, with a genomics-centered core and adjacent coverage in other ‘omics and quantitative biology settings. Each problem comprises an encapsulated multi-step analysis with staged data, prompts that define a quantity of interest while otherwise providing minimal guidance, and verifiable answers. Solving each problem requires identifying and addressing realistic obstacles such as measurement error, selection bias, confounding, QC failures, and choosing among competing model classes. Through extensive ablation studies, we verify that each problem admits a single defensible answer. Each problem involves multiple dependent decision points; *i.e*., substantive inferential forks where a plausible wrong choice changes the downstream analysis, such that errors propagate through the inferential chain and into the final graded target. In initial evaluations, the mainline GPT family reaches an eval-level pass rate of 25.0% with GPT-5.5 at the xhigh reasoning setting. In separately reported GPT Pro runs, GPT-5.5 Pro reaches 33.2%, GPT-5.4 Pro reaches 25.6%, and GPT-5.2 Pro reaches 10.8%. Even for the two strongest reported Pro-harness settings, 60.2% and 62.1% of problems, respectively, remain below 20% pass rate over repeated runs. The strongest external baseline, Gemini 3.1 Pro, achieves 11.2%. Models often complete substantial portions of the workflow but exhibit a consistent gap between *noticing* and *acting*: they identify local diagnostic signals but fail to propagate the implication to the corresponding analysis decision, and as a result select wrong estimators or persist on initially plausible but incorrect analysis paths. GeneBench therefore measures an emerging capability that remains as yet unreliable.

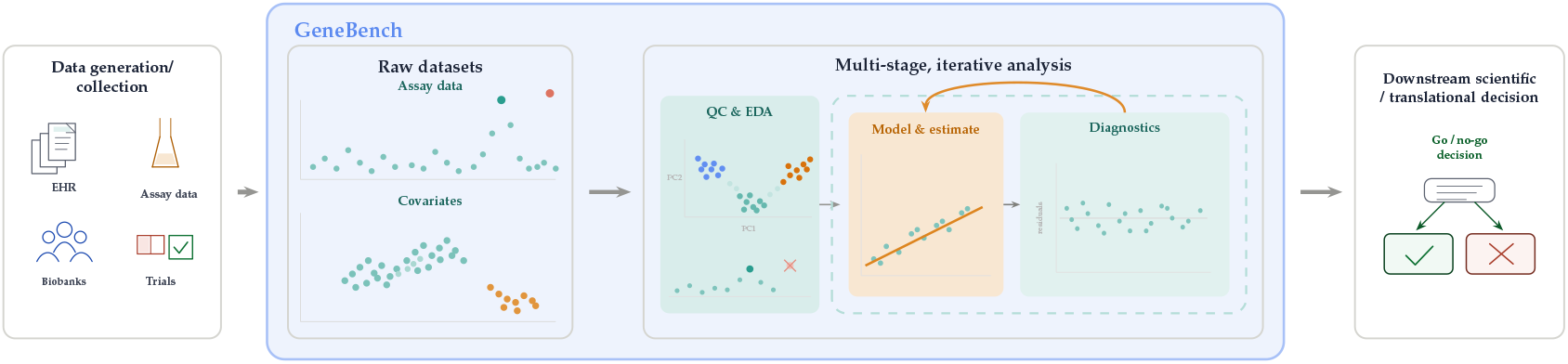

## Introduction

AI capabilities are advancing rapidly along multiple separate axes. Agentic systems now perform strongly on software engineering benchmarks such as SWE-Bench, SWE-Bench Verified, and SWE-Lancer; ^1,3,4^ broader evaluation efforts such as FrontierScience, FrontierMath, and Humanity’s Last Exam show similar progress on difficult expert-level and novel-problem settings, ^6–8^ and METR’s recent time-horizon analyses likewise suggest that the duration of tasks frontier agents can complete autonomously is increasing rapidly. ^9^ Simultaneously, biology foundation models such as ESM3 and Evo 2 have pushed protein and genome modeling to new scale and fidelity. ^56,57^

Yet there has been relatively little formal examination of AI performance on the broader routine process that underpins much of modern science: executing a quantitative analysis starting from potentially errorful raw data obtained from assays, clinical systems, or other data collection pipelines, and ending at a decision-relevant conclusion (see **Figure 1B** for a high-level schematic of the typical process). This class of work is a major practical bottleneck in data-rich fields including genomics, proteomics, transcriptomics, and metabolomics; for example, recent reviews in the genomics literature argue that as sequencing has scaled, downstream computation and analysis, rather than data generation, have become the central bottleneck. ^66–68^

**Figure 1:**
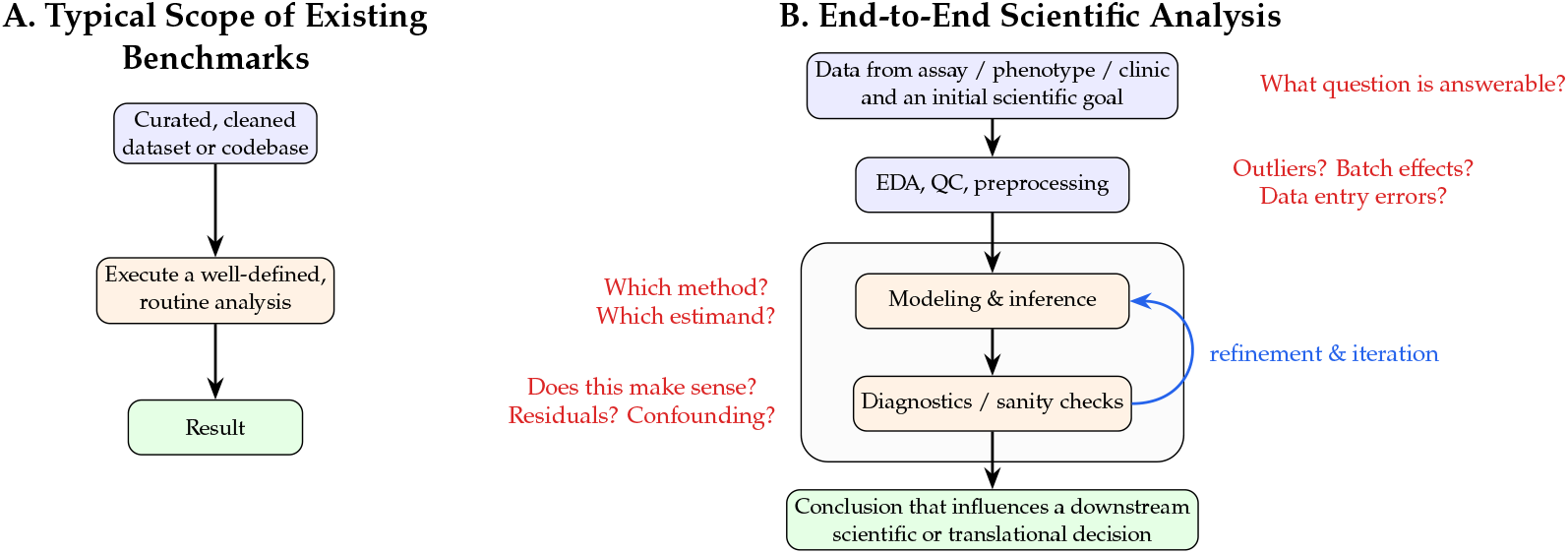
The benchmark gap. **(A)**: many current benchmarks in scientific AI begin from a curated dataset, codebase, or localized question and evaluate a narrowly scoped analysis step with a clearly verifiable answer. **(B)**: real-world scientific analysis more often spans a wider and more iterative process: data are obtained from an assay, clinic, or experiment; analysts must decide what question is answerable, perform quality control and exploratory analysis, choose models and estimands, diagnose failures, and ultimately reach a conclusion that can influence the next scientific or translational decision. GeneBench is intended to evaluate this broader workflow rather than only isolated substeps.

In contrast to most engineering tasks, scientific research is far more iterative, open-ended, and ambiguous. Its core challenges stem not from the execution of analytical workflows, but from the importance of scientific intuition or “research taste”: chains of judgment calls about what question the data can support, what data to include, which estimand or model is appropriate, whether diagnostics invalidate initial hypotheses, and when the evidence is strong enough to support a conclusion.

Recent biology benchmarks come closer to this target, but still mostly cover narrower forms of the workflow (see **Figure 1A**). For example, GeneTuring emphasizes genomic knowledge retrieval; ^60^ LAB-Bench and LABBench2 focus on practical biology research capabilities such as literature reasoning and database navigation, while BixBench, BAISBench, SpatialBench, scBench, and CompBioBench evaluate more realistic but still narrowly scoped computational biology and omics analyses. ^58,59,61–65^ Such benchmarks are also increasingly saturated, rendering them less useful for tracking model improvements.

Genomics is a rapidly growing, data-rich field where AI capabilities are increasingly scientifically and economically relevant. Since the first wave of large-scale GWAS, exemplified by the Wellcome Trust Case Control Consortium seven-disease study in 2007, ^36^ human genetics has become a major engine for mechanism discovery and target prioritization in biomedicine. ^38^ Sequencing costs have fallen dramatically, outpacing Moore’s law after the transition to next-generation sequencing, while the amount of sequence data produced continues to strain downstream computation and analysis. ^52,53^ At the same time, the field now operates on cohorts such as the UK Biobank and All of Us which contain linked molecular, phenotype, and health record data at a scale that would have been impossible a decade ago. ^54,55^ Human genetic evidence has also become one of the strongest empirical priors in drug discovery: classic analyses have demonstrated that drug mechanisms with genetic support are roughly twice as likely to lead to approved indications, ^39^ and updated estimates place the advantage at 2.6-fold. ^40^

This combination makes genomics an ideal domain for investigation into AI-driven scientific analysis: it is scientifically central, fast growing, and rich in realistic tasks where iterative analysis and chains of judgment calls matter. At the same time, many genomics problems remain benchmarkable because the data are structured, intended solutions can often be staged to yield identifiable targets, and plausible but incorrect analyses can be explicitly invalidated.

Here we introduce GeneBench, a novel benchmark spanning industry and academic-relevant subdomains of genomics as well as adjacent ‘omics and quantitative biology topics. Each problem is a self-contained, multi-step analysis that provides (1) a realistic, messy dataset intended to reflect the data a scientist would receive from a lab, EHR system, or other collection pipeline, and (2) a target estimand. That estimand is chosen to reflect a quantity that would inform a downstream decision in practice. The current suite contains 103 evaluations across 10 domains, with a genetics-centered core in population genetics, statistical genetics, quantitative genetics, and functional genomics, and adjacent coverage in spatial transcriptomics, cancer genomics, proteomics, clinical genetics, forensic genetics, and epigenomics.

In early evaluations, the mainline GPT family reaches 25.0% pass rate with GPT-5.5 at the xhigh reasoning setting. Separately reported GPT Pro runs reach 33.2% for GPT-5.5 Pro, 25.6% for GPT-5.4 Pro ^16^, and 10.8% for GPT-5.2 Pro. At the highest mapped setting available for each mainline model, pass rate rises from 3.5% for GPT-5^13^ to 9.4% for GPT-5.2^14^, 19.0% for GPT-5.4^15^, and 25.0% for GPT-5.5. Manual examination of the model-reported reasoning from GPT-5.4 and GPT-5.5 suggests that the main qualitative improvement in performance observed in stronger models is less in identifying and recognizing the relevant diagnostic clues than in turning such observations into concrete decisions on what corrective actions or model-selection decisions to take that move the analysis onto the correct analysis path.

In the remainder of the paper, we first describe the scope of GeneBench using a high-level atlas of the problem space, introduce the main design constraints required to make this class of decision-heavy scientific analysis benchmarkable, and review a representative example problem to make these abstractions concrete. We then present benchmark-wide results and discuss qualitative improvements in model performance. A detailed case study involving a genome-wide association study (GWAS) is provided in the **Appendix**.

## Benchmark Scope and Construction

GeneBench, a collection of 103 problems across 10 domains, measures whether an agent can recover a valid quantitative analysis from potentially errorful datasets with minimal guidance. Across the current suite, an agent must filter and correct data, identify QC or ascertainment problems, choose methods, revise the analysis when intermediate results disagree with the initial plan/hypothesis, and produce a final quantitative answer. Many problems are framed as decision points in genetics-backed drug discovery and translational research, such as whether a GWAS signal survives correction strongly enough to advance and which gene or protein should be nominated as the likely effector target, while others are framed around more academically oriented questions, such as whether an observed pattern is better explained by selection or demography and which pedigree, haplotype, or ancestry reconstruction is supported by the data. **Figure 2** illustrates the domain coverage of the current suite, and summaries of 23 representative problems are provided in **Supplementary Table 1**.

**Figure 2:**
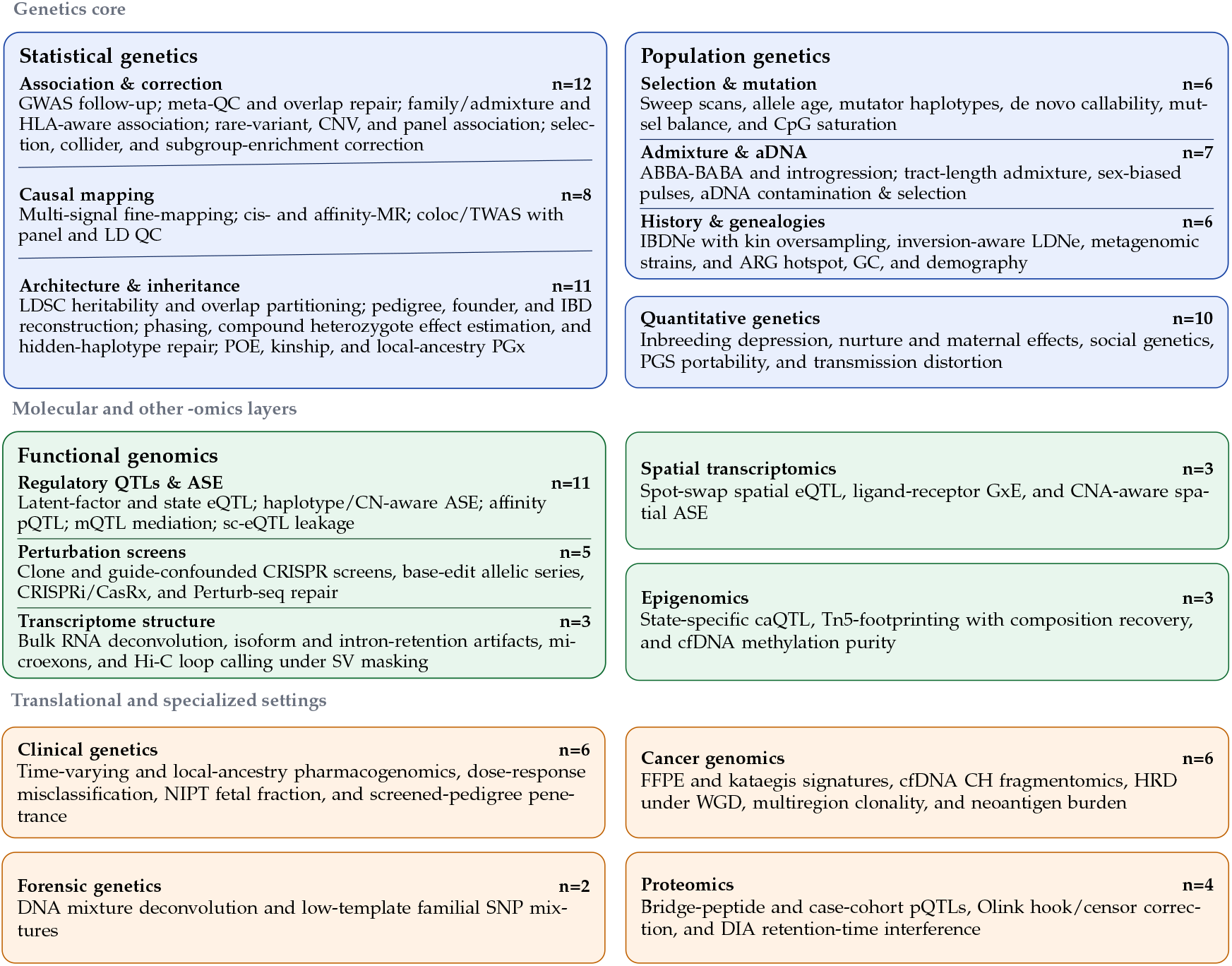
Domain atlas of the current GeneBench suite. GeneBench comprises 103 problems across 10 domains. Nested subcards expose the main subdomains within statistical genetics, population genetics, and functional genomics. *Abbreviations:* GWAS, genome-wide association study; QC, quality control; HLA, human leukocyte antigen; CNV, copy-number variant; MR, Mendelian randomization; TWAS, transcriptome-wide association study; LD, linkage disequilibrium; LDSC, linkage disequilibrium score regression; IBD, identity by descent; comphet, compound heterozygosity; POE, parent-of-origin effect; PGx, pharmacogenomics; CpG, cytosine-phosphate-guanine; aDNA, ancient DNA; IBDNe and LDNe, effective population-size inference from identity-by-descent and linkage disequilibrium, respectively; ARG, ancestral recombination graph; GC, gene conversion; PGS, polygenic score; ASE, allele-specific expression; eQTL, expression quantitative trait locus; pQTL, protein quantitative trait locus; mQTL, methylation quantitative trait locus; caQTL, chromatin-accessibility quantitative trait locus; sc-eQTL, single-cell expression quantitative trait locus; CRISPRi, CRISPR interference; CasRx, an RNA-targeting CRISPR effector; Hi-C, genome-wide chromosome conformation capture; GxE, gene-by-environment interaction; CNA, copy-number alteration; NIPT, noninvasive prenatal testing; cfDNA, cell-free DNA; FFPE, formalin-fixed, paraffin-embedded; CH, clonal hematopoiesis; HRD, homologous recombination deficiency; WGD, whole-genome doubling; DIA, data-independent acquisition.

### Benchmark Setup

Each GeneBench problem is packaged as a self-contained scientific analysis. The agent receives an isolated workspace containing a *minimum viable prompt*, staged files, and a standard scientific Python stack. The prompt specifies the scientific question/task and target estimand without explicitly prescribing the workflow to be executed. The files are intended to resemble what an analyst might actually receive from assays or clinical systems rather than cleaned toy datasets. Each problem involves a chain of dependent decision points such that an incorrect choice at any stage propagates into downstream errors and ultimately failure to recover the final correct target.

The sandbox in which the agent operates is relatively sparse, with the agent receiving only the staged files and access to general-purpose scientific libraries including `numpy, pandas, scipy, scikit-learn, statsmodels, lifelines, matplotlib`, and `seaborn`, but no domain-specific bioinformatics tooling or packages. **Supplementary Figure 1** shows a schematic of the agent environment. Success therefore depends both on the agent recovering the analysis from the data as well as accurate implementation of the relevant methods.

### Construction, Validation, and Grading

Open-ended scientific analysis is difficult to benchmark precisely because real data often admit multiple defensible analysis choices. For example, QC thresholds, model parameterizations, and reporting conventions can vary across analysts without there being only a single unambiguously correct analytical choice. If the outcomes of a benchmark change because one agent uses one defensible cutoff or convention while another agent uses a different, yet equally defensible one, this might reflect the arbitrary nature of that benchmark’s design choices rather than the quality of scientific reasoning.

Furthermore, real analyses involve multiple stages. For example, assays must be calibrated before association testing, and ascertainment biases must be corrected prior to effect estimation. Failure to execute any one stage can result in significant downstream changes in the analysis pipeline, and ultimately estimation of the final quantity of interest upon which a significant business or clinical decision depends. A useful benchmark for this type of work must therefore be insensitive to nearby defensible analyst choices, but sensitive to missing scientifically necessary stages. To model the multi-stage nature of realistic scientific workflows, GeneBench problems are intentionally “cascaded”, such that upstream decisions change what analyses are valid downstream. We quantify this cascaded structure through the number of *decision points* in each problem: substantive inferential forks where a plausible wrong choice leads to a qualitatively different downstream answer. The number of these decision points ranges from 3 to 13 across the current suite (with a median of 6), and are shown for the representative problems in **Supplementary Table 1**.

We count a decision point only when the staged data create a distinct inferential fork that a careful analyst could resolve from agent-visible evidence, and where a plausible wrong choice propagates into a materially different downstream analysis or graded answer. We do not count purely mechanical file handling or minor parameter tuning within an otherwise fixed method. Closely linked checks that jointly implement a single correction are also counted as one decision point rather than several. In order to implement these multi-stage setups, GeneBench problems rely on constructively simulated problems rather than historical real datasets. This lets us directly tune the number and difficulty of decision points while ensuring that (1) QC-sensitive decisions are robust to small researcher-choice variation, (2) plausible wrong analyses fail for substantive reasons, and (3) the graded endpoint is actually recoverable from the agent-visible data.

Operationally, problem development begins from a real-world analysis pattern and a target estimand. These real-world analysis patterns are synthesized from the literature and internal expertise to reflect common, high-impact scientific questions and workflows, and are specifically chosen so they do not recapitulate well-known textbook examples or papers, so as to avoid the risk of benchmarking against memorized solutions. Data are then simulated so that the correct answer is recoverable from the staged files (for example, the maximum likelihood estimate of a parameter resulting from the correct approach would be considered as the ground truth value for grading, rather than the parameter under which the data were generated). A minimum viable prompt containing the minimum amount of information required to make the correct answer identifiable is then constructed.

Once an initial draft of a problem is completed, extensive validation is performed. Results from analyses involving plausible but incorrect decisions at the various inference stages are checked via ablation and verified to be sufficiently distinct from the graded answer. Independent reviews for scientific validity, methodological soundness, and target identifiability are conducted in order to ensure that the evaluation is testing the intended capabilities rather than whether agents can guess the benchmark designer’s preferred (but non-unique) workflow. Problem drafts are then iteratively audited through multiple rounds of frontier-model pilots and detailed trace analyses in order to check for unintended leakage, alternative unintended pathways to the correct answer, prompt-grader mismatch, and robustness. This process is intended to ensure that wrong-but-plausible analyses fail for substantive reasons and that passing runs reflect the intended inferential path rather than shortcuts. **Table 1** summarizes the main benchmark-level constraints that follow from these requirements.

**Table 1.** Primary design constraints in GeneBench. Together, these are intended to keep the graded endpoint scientifically identifiable while preserving realistic ambiguity, data error, and multi-stage analysis complexity.

| Principle | Benchmark requirement | Failure mode if violated |
| --- | --- | --- |
| *Ground truth and identifiability* |  |  |
| Recoverable target | Agents are graded on recovering the quantity that is actually recoverable from agent-visible data, and not the hidden data-generating parameters. | Correct analyses can be marked wrong because the parameter under which data were generated is unrecoverable (*e.g.*, due to sampling variation in the DGP). |
| Unique, identifiable answer | The staged evidence along with a minimum viable prompt supports one uniquely defensible answer. If multiple approaches would ordinarily appear defensible, the data contain some empirical signature that rules out all but one. | The task becomes under-specified, and success depends on guessing the benchmark designer’s preferred pipeline rather than reasoning from the evidence. |
| Clear separation from incorrect answers | A comprehensive ablation suite demonstrates that plausible wrong analyses and shortcut methods yield materially different answers and fail by clear margins. | Wrong analyses land too close to the target, so grading depends on tolerance tuning rather than scientific correctness. |
| *Problem specification* |  |  |
| Minimal viable prompt | The scientific question, graded estimands, conventions, and output format are defined clearly on the agent-visible surface, but the prompt does not hint at the method, QC path, or intermediate workflow. | The task either collapses into prompt following or leaves multiple defensible interpretations of what answer should be reported. |
| Threshold-robust QC | When QC is part of the solution, nearby reasonable thresholds lead to the same graded outcome. | The benchmark measures arbitrary cutoff choice rather than recognition of the QC problem. |
| *Scientific workflow fidelity* |  |  |
| Constructive staging | Simulating data allows us to tune each detail of the data-generating process so realism, multi-stage inference, effect sizes, and diagnostic clues can be precisely controlled. | Difficulty, answer separation, and correctness become difficult to calibrate. |
| Multi-stage inference | Upstream filtering, representation, and adjustment-model decisions materially affect the final graded endpoint. | The benchmark reflects smaller units of end-to-end analysis rather than the full flow. |
| Literature-defensible solution | The intended correct solution involves standard or otherwise well-supported methods. | Success depends on benchmark-specific machinery, ad-hoc designer choices, rather than scientific judgment. |

At present, GeneBench uses binary grading against recoverable targets under calibrated tolerances chosen to allow for numerical and implementation-level variation; the evaluation setup and package-level grading protocol is summarized in the **Methods**.

### Example Problem: LDL-C GWAS Follow-up

**Figure 3** shows a representative GeneBench problem from statistical genetics. The task is framed as follow-up on a candidate signal from a genome-wide association study (GWAS) of low-density lipoprotein cholesterol (LDL-C), a routine cardiovascular biomarker. In practice, analyses of this kind are used to determine whether an apparent association survives standard QC and design corrections. ^47,48^ They are then used to determine whether the signal is interpretable enough to motivate downstream biological follow-up. ^38^

**Figure 3:**
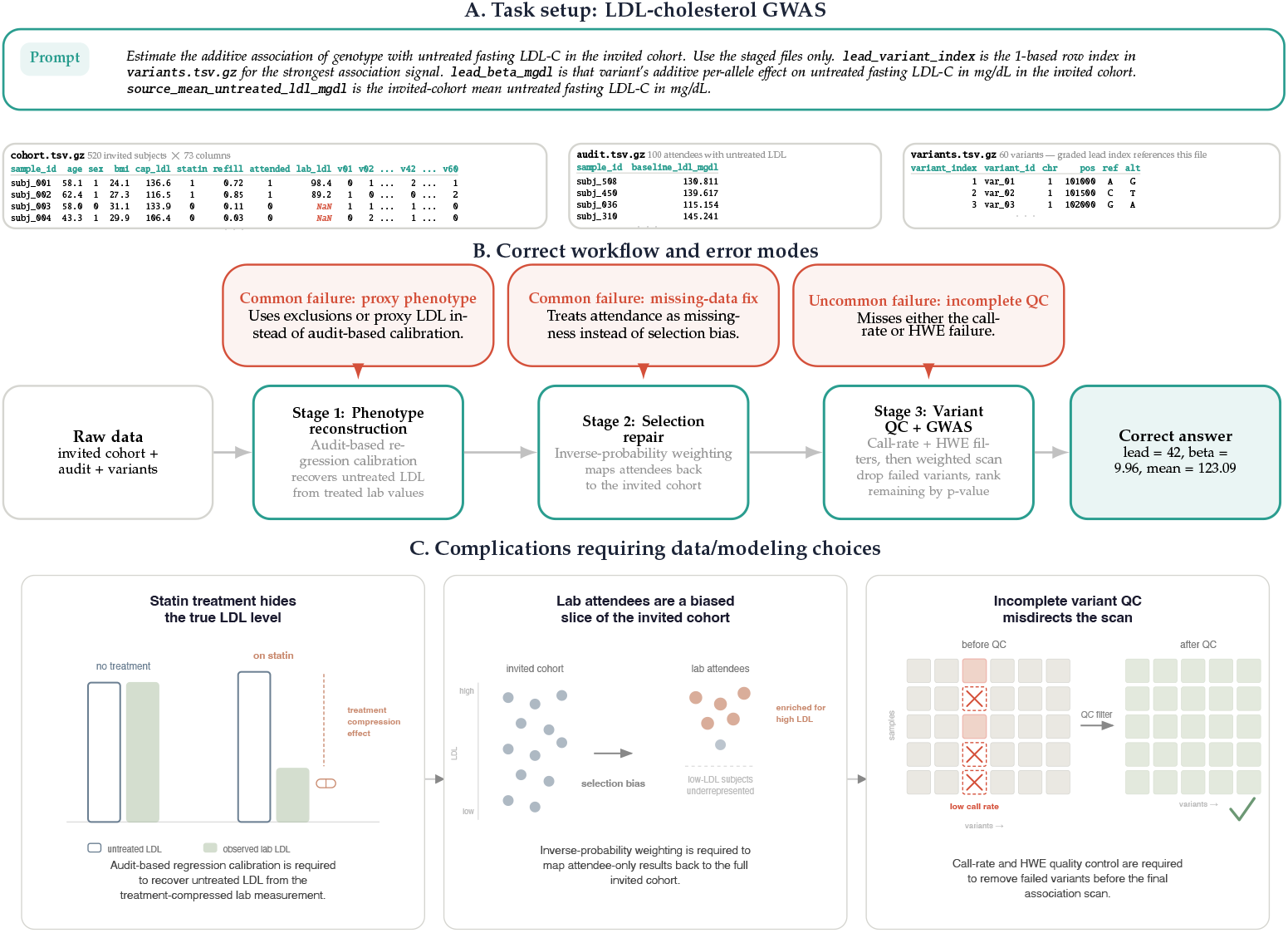
Representative GeneBench problem from statistical genetics: LDL-cholesterol GWAS follow-up. **(A)** Problem setup. The agent receives a sparse prompt and three staged files: a cohort table for the full invited cohort, an audit subset with untreated LDL-C, and a variant manifest. The graded target is the additive association of genotype with untreated LDL-C in the invited cohort. **(B)** Correct analysis path and representative failure modes. Recovering the target requires phenotype reconstruction from the audit data, reweighting attendees back to the invited cohort, and variant QC before the final association scan. The red nodes mark shortcut analyses seen in model traces and included in the ablation studies: local repairs that address one visible problem but stop short of the full inferential chain required to recover the ground-truth result. **(C)** Decision points within the inferential chain. Each stage addresses a different distortion in the staged data: treatment masks the phenotype, attendance is selective, and two variants carry distinct technical QC failures requiring both call-rate and Hardy–Weinberg filtering. Upstream choices therefore determine which downstream analyses remain valid. The full prompt, construction, and ablation evidence are given in the **Appendix**.

Here the agent receives a sparse prompt, and three data files: (1) a table with phenotypes/covariates for an invited cohort, (2) a small audit dataset, and (3) a manifest of genetic variants. The staged data are structured around three nested views of the cohort. The cohort table contains the full invited cohort together with covariates, a noisy capillary LDL-C proxy measured at invitation, statin treatment status, prescription refill data, and fasting-lab LDL-C for the subset who later returned for a standardized clinic blood draw after fasting. This fasting-lab measurement is the cleaner clinical phenotype, but it is only observed for attendees. The audit subset is a random subset of those attendees for whom a historical untreated LDL-C measurement is also available. The variant manifest provides the candidate variants to be screened in the final association scan.

The graded target is the additive association of genotype with *untreated fasting LDL-C in the invited cohort*, rather than with the *observed LDL-C values among the subset who returned for that fasting visit*. This distinction matters because statin treatment in the cohort compresses the measured phenotype relative to the untreated state, and those who attended the fasting visit are a selected subset. Additionally, one candidate variant in the manifest carries a genotype-missingness artifact, while another fails a basic Hardy–Weinberg equilibrium check. The task therefore cannot be solved by a naive association scan on the returned lab values. Valid inference requires recognizing the need for reconstructing untreated LDL-C from the audit data, reweighting attendees back to the invited cohort, and applying both call-rate and Hardy–Weinberg variant QC before the final scan.

Each stage reflects a decision point for the agent, which must reason through the implications of each data feature, choose whether to execute any sort of corrective action and of what kind, and follow through on the correct analysis.

## Results

We evaluated a selection of models on the full 103-problem GeneBench suite, comparing recent models in the GPT family with external non-GPT baselines. Models in the mainline GPT family tested were GPT-5^13^, GPT-5.2^14^, GPT-5.4^15^, and GPT-5.5 across the reasoning-effort settings available for each model. We also report results from GPT-5 Pro ^17^, GPT-5.2 Pro ^18^, GPT-5.4 Pro ^16^, and GPT-5.5 Pro. External models tested were MiMo V2 Pro ^25^, MiMo V2.5 Pro ^26^, Kimi K2.5^23^, Kimi K2.6^24^, Grok 4.20^20^ with reasoning enabled, Qwen 3.6 Plus ^21^, GLM 5.1^22^ with reasoning enabled, and Gemini 3.1 Pro ^19^. We do not report results for models from Anthropic due to Terms of Service restrictions. **Figure 4** summarizes the overall pass rates across the suite.

**Figure 4:**
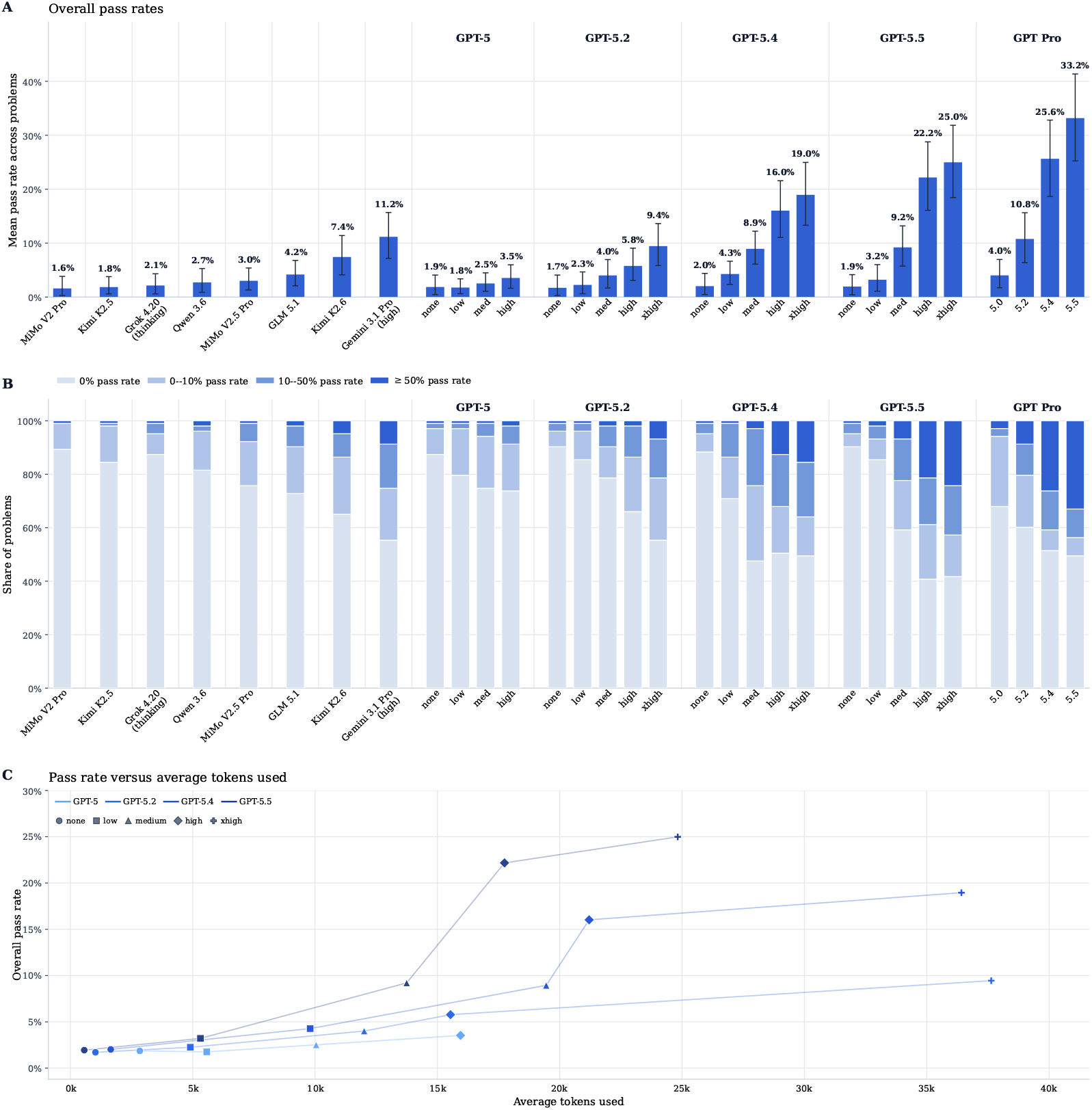
Benchmark-wide performance across evaluated model settings. **(A)** Overall pass rate, defined as the unweighted mean of per-problem pass rates across the 103 benchmark problems. Error bars show 95% hierarchical bootstrap confidence intervals from 20,000 resamples, obtained by resampling problems and, within each sampled problem, run-level outcomes. **(B)** Distribution of per-problem pass rates across four regimes: 0%, 0–10%, 10–50%, and at least 50%. **(C)** Overall pass rate versus average tokens used per problem for the GPT family. Average tokens used per problem was computed as the number of tokens in the model’s full chain-of-thought trace and final response, excluding tool calls. Line colors denote model families, and point shapes denote effort settings. The four rightmost bars in panels A and B correspond to separately reported Pro-harness runs. GPT-5 Pro, GPT-5.2 Pro, GPT-5.4 Pro, and GPT-5.5 Pro are omitted from panel C. GPT-5 is shown from none through high, and later mainline GPT models are shown from none through xhigh. Gemini 3.1 Pro is shown with high reasoning effort, and Grok 4.20 is shown with reasoning enabled but no explicit reasoning-effort tier.

For each model, we ran multiple replicates of each of the 103 problems. Across reported model-problem pass rates, the number of runs (see **Methods**) averaged 28.7 and ranged from 14 to 60 depending on the model setting. This variation affects precision, and therefore the width of the confidence intervals, but not the point estimates in **Figure 4A**, which are computed as unweighted means of per-problem pass rates rather than pooled pass rates over all runs.

### Overall performance and unsolved tail

Outside the separately reported Pro-harness runs, overall pass rates range from 1.6% for MiMo V2 Pro to 25.0% for GPT-5.5 at the `xhigh` reasoning setting. At the matched `xhigh` reasoning setting within the GPT family, mean pass rate rises from 9.4% for GPT-5.2 to 19.0% for GPT-5.4 and 25.0% for GPT-5.5. GPT-5 reaches 3.5% at its highest mapped setting, `high`. Among external models, Gemini 3.1 Pro reaches 11.2%, exceeding GPT-5.2 at the `xhigh` reasoning setting and GPT-5.2 Pro. Kimi K2.6 reaches 7.4%, below Gemini but above the other external baselines. MiMo V2.5 Pro reaches 3.0%, above MiMo V2 Pro but below GLM 5.1. GLM 5.1 reaches 4.2%, exceeding GPT-5 at all reported effort settings but remaining below Gemini 3.1 Pro and Kimi K2.6. The Pro-harness runs, reported separately under that special-case setup, reach 4.0% for GPT-5 Pro, 10.8% for GPT-5.2 Pro, 25.6% for GPT-5.4 Pro, and 33.2% for GPT-5.5 Pro. Within each later GPT model family, reasoning effort is a major determinant of performance: pass rate rises from approximately 2% at `none` to 9.4%, 19.0%, and 25.0% at `xhigh` for GPT-5.2, GPT-5.4, and GPT-5.5, respectively (**Figure 4C**).

A substantial unsolved tail remains (**Figure 4B**). Along the mainline GPT progression at the highest mapped setting for each model, the share of problems with 0% pass rate declines from 73.8% to 55.3% to 49.5% to 41.7%, whereas the share reaching at least 50% rises from 1.9% to 6.8% to 15.5% to 24.3%. The benchmark therefore remains dominated by more difficult items, although stronger models move a larger fraction of problems out of the floor regime and into partial or frequent success, consistent with broad-based increases in model intelligence. Exact values underlying **Figure 4**, external-model token counts, and the corresponding numbers of runs per problem are reported in **Supplementary Table 2**. The separately reported Pro-harness runs also remain far from saturation: GPT-5.2 Pro, GPT-5.4 Pro, and GPT-5.5 Pro leave 87.4%, 62.1%, and 60.2% of problems below 20% pass rate, respectively, while 8.7%, 26.2%, and 33.0% of problems reach the ≥ 50% regime.

### Inferential chain length and action on intermediate diagnostic evidence

Pass rate declines with the length of the required inferential chain, measured here as the number of decision points in each problem (**Supplementary Figure 2A**). Problems with shorter chains (3–4 decision points) are solved at materially higher rates than those with longer chains (7+), and this gradient is steepest for the strongest models (**Supplementary Figure 2B**). Weaker models remain near the floor regardless of chain length, whereas stronger models can often solve shorter chains but break down as the number of required correct sequential inferences increases.

Manual review of the model-reported reasoning from GPT-5.4 and GPT-5.5 suggests a consistent mechanism behind this scaling. In most failures, the agent notices the relevant local clue/diagnostic but does not propagate the conclusion it ought to make from that into the relevant downstream analysis decision. **Table 2** shows representative excerpts. For example, the weaker model, GPT-5.4, often applies a partial QC fix, over-corrects with redundant ancestry covariates, or makes a local sign correction without carrying it through to the final reported answer, whereas the stronger model, GPT-5.5, is more likely to carry the same diagnosis through to the final analytical choice.

**Table 2:** Representative excerpts from the model-reported reasoning of selected GPT-5.4 and GPT-5.5 comparisons. In each case, both models identify or note the relevant local signal, but the stronger model carries the diagnostic through to a corresponding change in the downstream analytical choice.

| Problem | GPT-5.4 | GPT-5.5 |
| --- | --- | --- |
| **Somatic signature deconvolution with FFPE and kataegis:** FFPE strand bias and localized kataegis bursts both inflate APOBEC (SBS2/SBS13) exposures and must be filtered before per-sample signature deconvolution. | **Tests removing kataegis, observes the estimate change, then deliberately but incorrectly retains them.**<br>"Removing very clustered kataegis-like mutations reduced $\beta$ to about 0.72, indicating that some of the association is carried by clustered APOBEC events; because the target parameter is the total SBS2+SBS13 fraction, I retained those clustered mutations in the primary estimate." | **Removes kataegis clusters explicitly before deconvolution.**<br>"used a high-confidence mutation set . . . and removed localized kataegis/clustered bursts defined as $\geq 6$ same-sample same-chromosome mutations within 10 kb." |
| **SNP heritability of cortisol in a multi-ancestry cohort:** including ancestry-PC fixed effects beyond the true ancestry axes absorbs heritable signal and biases $h^2$ toward zero. | **Adds many ancestry PCs and reports a strongly attenuated $h^2$.**<br>"computed 20 genotype principal components from standardized dosages . . . included them as additional fixed covariates . . . The resulting SNP heritability estimate was $h^2 = \sigma_g^2 / (\sigma_g^2 + \sigma_e^2) = 0.122$." | **Uses two scree-elbow PCs and recovers a value near the truth.**<br>"the supplied PC1 matched the first genotype PC, and I included the first two genotype PCs as fixed effects based on the GRM eigenvalue elbow . . . REML . . . estimated $h^2 = \sigma_g^2 / (\sigma_g^2 + \sigma_e^2) = 0.403$ on the log-cortisol scale." |
| **Multi-signal fine-mapping with a sign-flipped palindromic variant:** rs100055 has a palindromic A/T allele pair and a sign-flipped summary effect; the agent must detect the LD/$z$ inconsistency, flip the effect, and carry the flipped direction through to the reported sign. | **Flips the sign for internal LD coherence but reports the original unflipped direction in the final answer.**<br>"applied the minimal sign correction needed for local $z$-LD coherence there before fine-mapping . . . Effect directions reported in the answer are from the original REF/ALT-aligned summary betas: ALT increases the trait for both rs100020 and rs100055." | **Carries the QC sign correction through to the final reported direction.**<br>"rs100055 had an LD-sign inconsistency with its high-LD neighbors; flipping its effect sign made the regional $z$-score pattern coherent, so I treated it as a strand/sign QC correction . . . the ALT allele T at rs100020 increases the trait, while the ALT allele T at rs100055 decreases the trait." |

On the most difficult tasks with the longest inferential chains where the pass rate remains 0% across all models, failures typically occur in this manner at one of the intermediate decision points: the model identifies the right warning sign but does not revise the analysis path enough to reach the valid final inference through the following steps. Taken together, these results suggest that GeneBench difficulty is driven by the intended linking of diagnostics to corrective action across a sequence of dependent decisions.

## Discussion

Agentic abilities in software engineering, computer use, broad scientific reasoning, and general capabilities have been increasing at a rapid pace, as evidenced by recent model progress on benchmarks evaluating these skills ^3–5,8,60,61^. However, the types of open-ended, multi-stage scientific analyses that are common to real-world research and industrial applications remain underexamined.

GeneBench is a new genetics and quantitative biology evaluation intended to target this gap. In our initial evaluations, the strongest models already show substantial partial competence across many tasks, even when they do not complete the full decision-making chain. We observe that while frontier models consistently notice data issues, statistical irregularities, and other potential problems, there remains an incomplete ability to bridge the “notice-act” gap required to close the inferential loop. Qualitatively, this pattern resembles expert-novice differences in scientific problem solving observed in humans, where experts utilize their experience to guide problem representation and adaptive decision-making, while novices struggle to integrate observations into the broader context of the problem. ^69,70^ We therefore anticipate that improvements in planning, self-revision, and uncertainty-aware control should translate into meaningful gains on this class of work. ^10–12^

Realizing these capability gains depends on having evaluations that can reliably measure progress; while GeneBench is a first step at evaluating this gap in capabilities, it is not without limitations. Constructive staging and simulation make the endpoint identifiable and the grading interpretable, but GeneBench does not attempt to reproduce the documentation gaps, data scale, and study-specific irregularities of true real-world analyses. ^51^

A deeper limitation, shared with most AI benchmarks, is that our binary pass/fail grading collapses the stage-level evidence that our review of the model-reported reasoning suggests is most diagnostic of model capability, treating a run that executes six of seven decision points correctly as indistinguishable from one that fails at the first step. Future versions of GeneBench may move toward rubric-based and stage-level scoring, drawing on the rapidly developing literature on process reward models and rubric-based supervision for multi-turn agents. ^27–29^

Stage-level scoring is also a prerequisite for using GeneBench-style problems as the substrate for the dense per-turn reward and credit-assignment methods that have emerged for long-horizon agentic RL. ^30,31,33^ The explicit decision-point decomposition turns each problem into a sequence of intermediate targets rather than a single terminal outcome, which a growing body of work identifies as a key ingredient for credit assignment in multi-turn agent trajectories where episode-level signal is otherwise too sparse to be informative. ^32,34^ The failure mode we observe, *i.e*., that models notice the relevant diagnostic but do not act on it, also aligns closely with the explicit target of recent self-correction RL methods. ^35^

Enabling agents to reliably automate this class of analysis could significantly accelerate scientific discovery. Human genetic evidence has played an increasingly central role in target prioritization and translational follow-up, ^38^ where mechanisms with human genetic support are materially more likely to translate into approved indications. ^2,39,40^ The plummeting costs of sequencing and the expansion of biobank-scale resources with linked molecular, phenotype, and health record data have enabled this trend to accelerate, but one of its consequences is that the bottleneck is increasingly shifting from data generation to the ability to turn data into actionable insights.

Models that could consistently execute the types of analyses that currently require teams of expert analysts would therefore have a transformative impact on the throughput and nature of industrial research by accelerating hypothesis triage, target follow-up, and the iteration cycle between data generation and decision-making. As a rough point of reference, executed unaided by a human expert, a typical GeneBench problem would take on the order of 10–40 hours all-in. At a conservative $100–$200 per hour, the human labor cost of a single problem is already on the order of a few thousand dollars. By comparison, at current frontier-model API rates (on the order of $10–$30 per million output tokens) and the tens of thousands of tokens typically consumed per attempt (**Supplementary Table 2**), a single model attempt costs well under a dollar, *i.e*., three to four orders of magnitude below the human baseline per attempt, and still two to three orders of magnitude below it after dividing by the observed per-attempt pass rate. These figures are only illustrative, but they indicate that the operational value of reliable automation on tasks of this type could be substantial even before considering the effects of scale or accelerated iteration speed.

Our results indicate that while current models have made substantial progress toward automating these analyses, there remains a significant capabilities gap that separates current frontier models from the reliable end-to-end performance required to fulfill this potential.

## Methods

### Evaluation and grading

Evaluation was conducted on the full 103-problem GeneBench suite. Across problem-model configurations reported in the main text, we collected a mean of 28.7 valid independent runs, with a range of 14 to 60. The evaluated models were MiMo V2 Pro, MiMo V2.5 Pro, Kimi K2.5, Kimi K2.6, Grok 4.20 with reasoning enabled, Qwen 3.6 Plus, GLM 5.1 with reasoning enabled, Gemini 3.1 Pro, GPT-5, GPT-5.2, GPT-5.4, GPT-5.5, GPT-5 Pro, GPT-5.2 Pro, GPT-5.4 Pro, and GPT-5.5 Pro. MiMo, Kimi, Grok, Qwen, and GLM were accessed through OpenRouter. Gemini 3.1 Pro was accessed directly through the Gemini API. GPT-family models were accessed through OpenAI’s internal API-like interface. For the mainline GPT-family models reported in the main text, **Supplementary Table 2** lists the reasoning effort for each row. The Pro variants were run under a separate Pro harness configuration and are therefore reported separately from the mainline progression. Mainline and external-baseline runs were provided access to the same harness within a Linux environment in a Docker container with Python and standard scientific computing libraries (`numpy, pandas, scipy, scikit-learn, statsmodels, lifelines, matplotlib`, and `seaborn`). The execution environment had no internet access; agents were limited to the prompt, staged files, installed software, and model-internal knowledge. Average tokens used was defined as the number of tokens in the model’s full chain-of-thought trace and final response, excluding tool calls.

Gemini 3.1 Pro was run through the Gemini API with `high` reasoning effort. The OpenRouter-routed settings were as follows. All OpenRouter-routed runs used `max_output_tokens=65536`. Grok 4.20 used `x-ai/grok-4.20` with `reasoning_enabled=True` and no explicit reasoning-effort tier. Qwen 3.6 Plus used `qwen/qwen3.6-plus:free` with no explicit reasoning setting. GLM 5.1 used `z-ai/glm-5.1` with `reasoning_enabled=True`. Kimi K2.5 used `moonshotai/kimi-k2.5` with `reasoning_enabled=True`. Kimi K2.6 used `moonshotai/kimi-k2.6` with `reasoning_enabled=True` and no explicit reasoning-effort tier. Xiaomi MiMo V2 Pro used `xiaomi/mimo-v2-pro` with `reasoning_enabled=True`. Xiaomi MiMo V2.5 Pro used `xiaomi/mimo-v2.5-pro` with `reasoning_enabled=True` and no explicit reasoning-effort tier.

For each problem, the model is supplied with a series of initial instructions in the following order:

- a brief system message describing the container execution environment,
- the content of the prompt specifying the question at hand,
- instructions to return the final answer in a prespecified JSON schema including both numerical estimates and a brief, free-form summarization of its reasoning, and
- an enumeration of the locally mounted locations of all relevant data files.

Decision-point counts were assigned during problem construction and validation under a fixed rubric. A candidate step counted only if it represented a distinct inferential fork that was resolvable from agent-visible evidence and for which a plausible wrong choice produced a materially different downstream analysis or graded answer. Routine bookkeeping, file-format handling, generic EDA, and small parameter adjustments within an otherwise fixed method were not counted. Tightly coupled checks serving a single scientific correction were collapsed into one decision point.

Binary grading was performed based on pre-specified problem-specific target fields, exact-match rules, and absolute numeric tolerances. A run is counted as passing only if all graded fields satisfied their respective constraints. We report pass rates over repeated runs as the primary benchmark metric and do not describe internal partial-credit or diagnostic scoring pathways here. A free-text reasoning field is also collected for qualitative analysis but is not graded. Model responses were automatically graded by Python scripts encoding these constraints.

A small minority of runs (fewer than 1%) with invalid execution traces due to container- or tooling-related failures were excluded from analysis. Models were not subject to an additional uniform wall-clock budget imposed by our harness; runs remained subject to provider and platform behavior in the evaluation stack.

## Acknowledgements

We thank Joseph Pickrell and Joy Jiao for helpful discussions and feedback on earlier drafts of this manuscript.

---

**Supplementary Figure 1:**
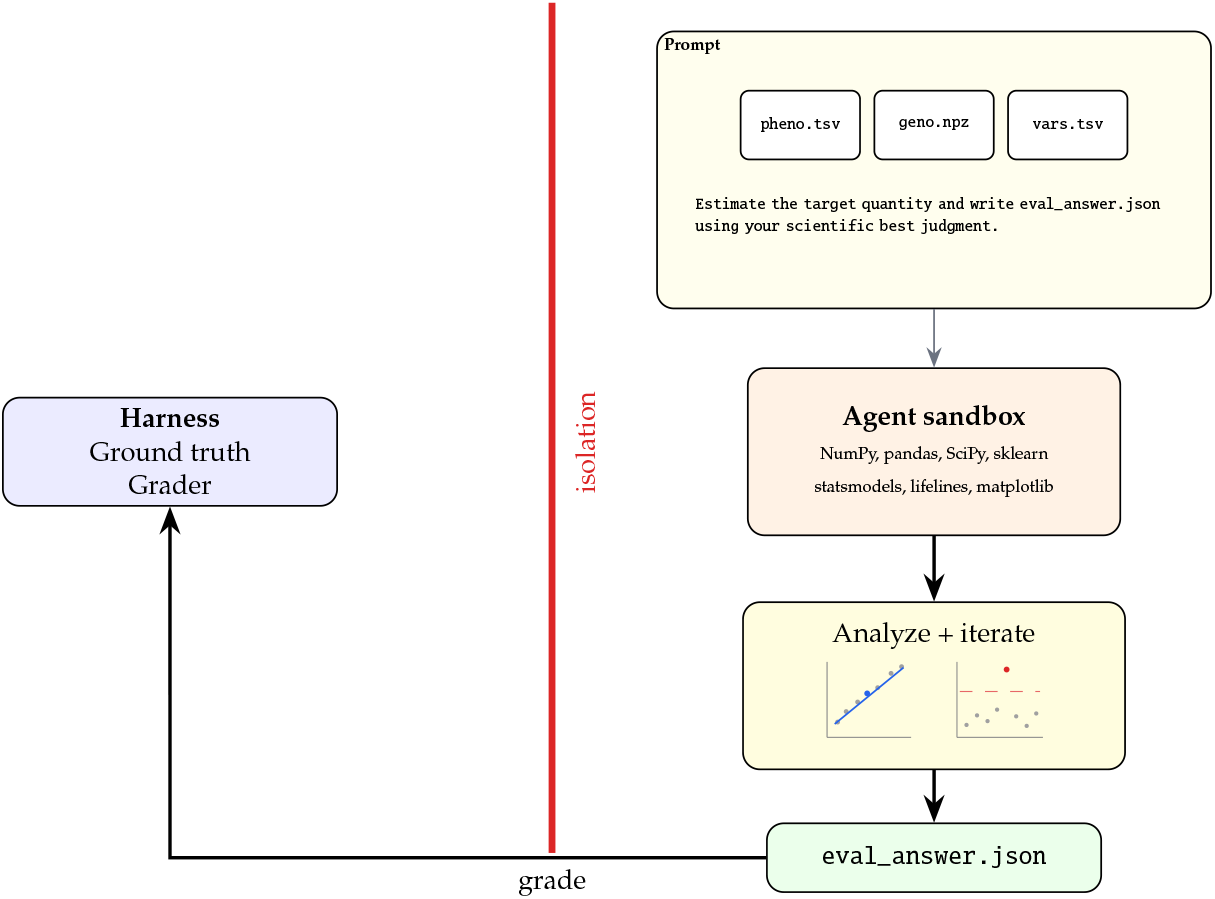
Agent environment and GeneBench problem anatomy. An agent receives a prompt, a set of files in an isolated workspace, and general-purpose scientific libraries. It must explore the data, test hypotheses, and execute an analysis before producing a final estimate of the target quantity in JSON format for grading.

**Supplementary Figure 2:**
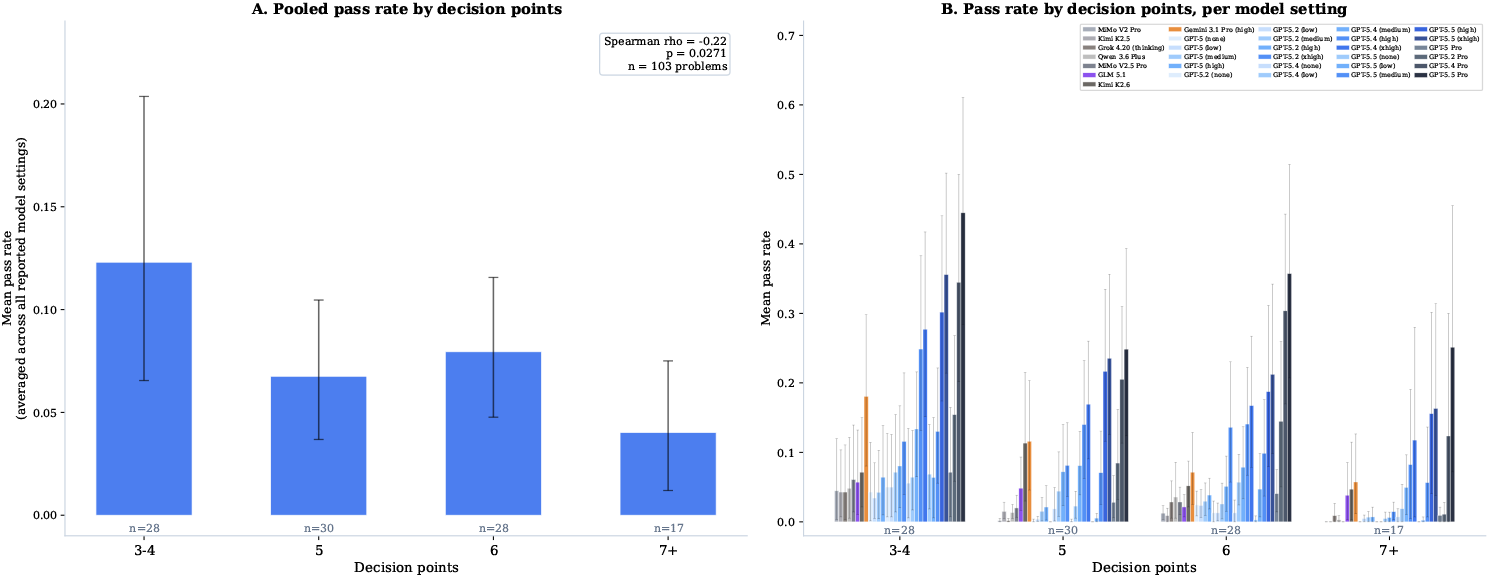
Pass rate by number of decision points. **(A)** Mean pass rate pooled across all reported model settings, binned by the number of decision points in each problem. Error bars show 95% bootstrap confidence intervals. **(B)** The same breakdown shown per model setting. Higher-performing settings still decline as the number of decision points increases.

**Supplementary Table 1:**
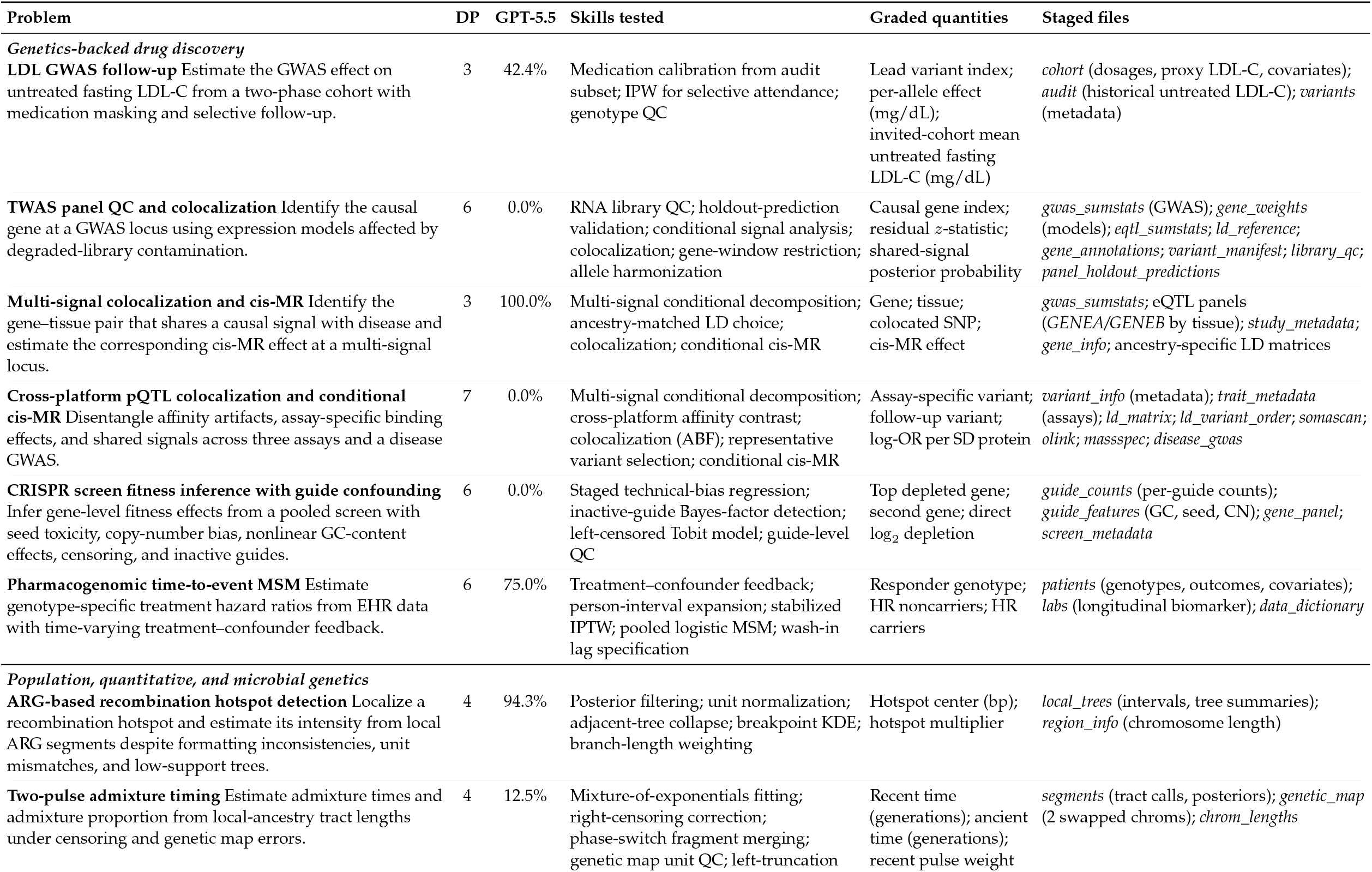
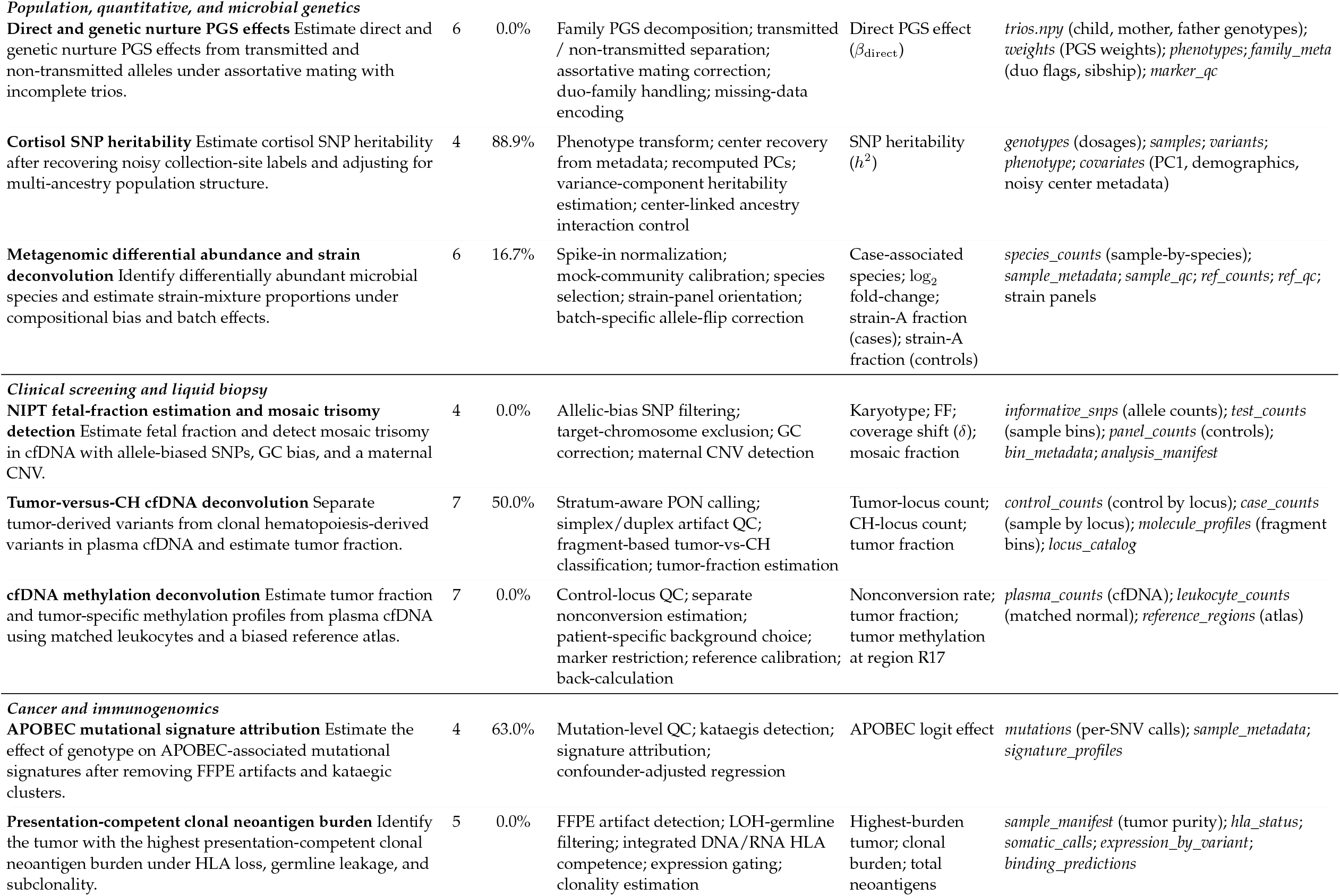
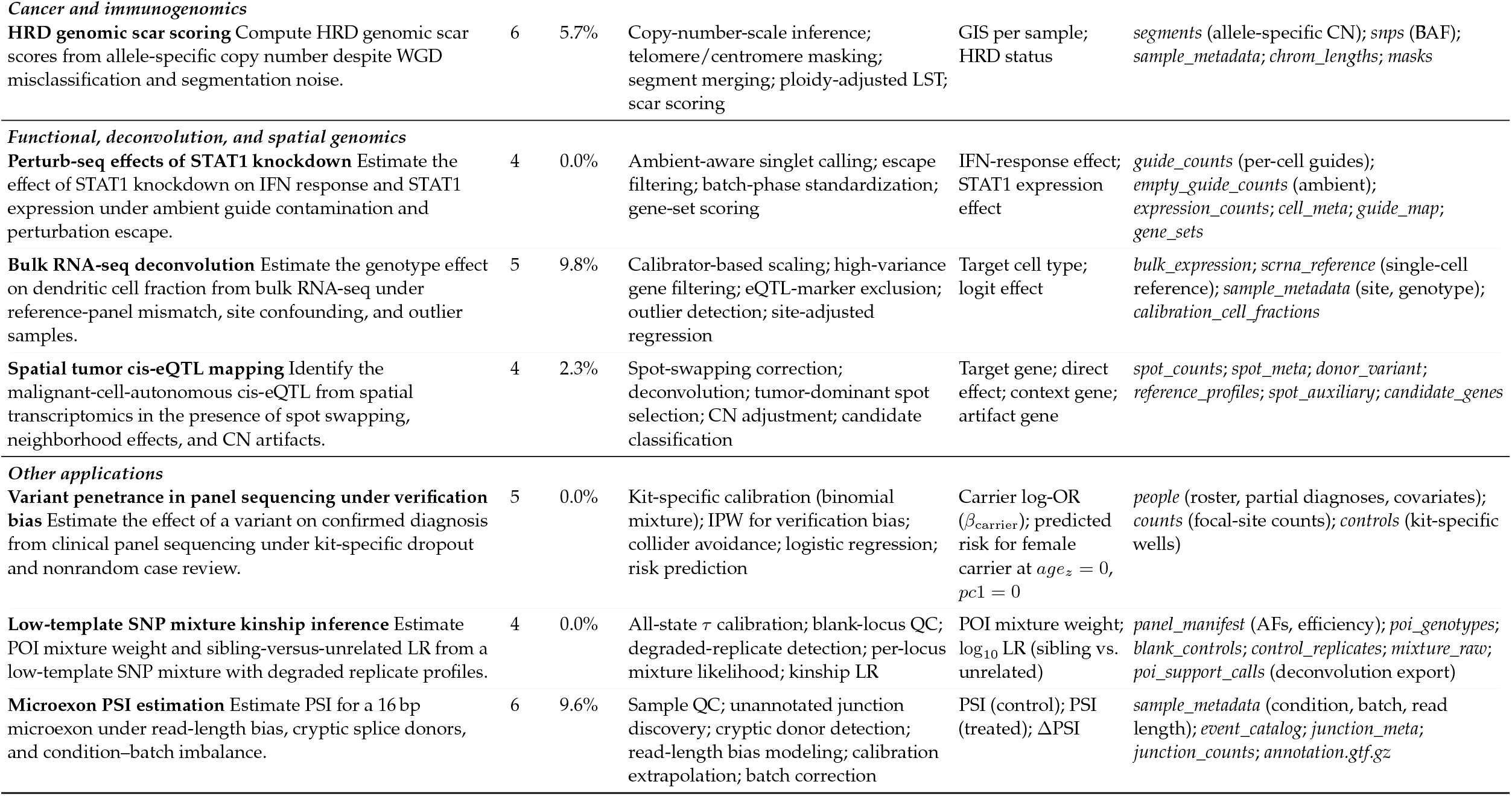
Twenty-three representative GeneBench problems. **DP** = decision points: substantive inferential forks where a plausible wrong choice leads to a qualitatively different answer. **GPT-5.5** reports pass rate for GPT-5.5 at the xhigh reasoning setting over repeated runs. *Abbreviations:* GWAS, genome-wide association study; LDL-C, low-density lipoprotein cholesterol; IPW, inverse-probability weighting; QC, quality control; TWAS, transcriptome-wide association study; coloc, colocalization; cis-MR, cis-Mendelian randomization; LD, linkage disequilibrium; SNP, single-nucleotide polymorphism; eQTL, expression quantitative trait locus; pQTL, protein quantitative trait locus; ABF, approximate Bayes factor; CN, copy number; GC, guanine-cytosine; PGx, pharmacogenomics; HR, hazard ratio; EHR, electronic health record; IPTW, inverse-probability-of-treatment weighting; MSM, marginal structural model; ARG, ancestral recombination graph; KDE, kernel density estimation; PGS, polygenic score; PC, principal component; NIPT, noninvasive prenatal testing; cfDNA, cell-free DNA; CNV, copy-number variant; FF, fetal fraction; PON, panel of normals; CH, clonal hematopoiesis; FFPE, formalin-fixed, paraffin-embedded; APOBEC, apolipoprotein B mRNA editing catalytic polypeptide-like; HLA, human leukocyte antigen; LOH, loss of heterozygosity; HRD, homologous recombination deficiency; WGD, whole-genome doubling; BAF, B-allele frequency; LST, large-scale state transition; GIS, genomic instability score; IFN, interferon; OR, odds ratio; POI, person of interest; LR, likelihood ratio; AF, allele frequency; SNV, single-nucleotide variant; PSI, percent spliced in; SD, standard deviation.

| Problem | DP | GPT-5.5 | Skills tested | Graded quantities | Staged files |
| --- | --- | --- | --- | --- | --- |
| **Genetics-backed drug discovery** |  |  |  |  |  |
| **LDL GWAS follow-up** Estimate the GWAS effect on untreated fasting LDL-C from a two-phase cohort with medication masking and selective follow-up. | 3 | 42.4% | Medication calibration from audit subset; IPW for selective attendance; genotype QC | Lead variant index; per-allele effect (mg/dL); invited-cohort mean untreated fasting LDL-C (mg/dL) | *cohort* (dosages, proxy LDL-C, covariates); *audit* (historical untreated LDL-C); *variants* (metadata) |
| **TWAS panel QC and colocalization** Identify the causal gene at a GWAS locus using expression models affected by degraded-library contamination. | 6 | 0.0% | RNA library QC; holdout-prediction validation; conditional signal analysis; colocalization; gene-window restriction; allele harmonization | Causal gene index; residual $z$-statistic; shared-signal posterior probability | *gwas_sumstats* (GWAS); *gene_weights* (models); *eqtl_sumstats*; *ld_reference*; *gene_annotations*; *variant_manifest*; *library_qc*; *panel_holdout_predictions* |
| **Multi-signal colocalization and cis-MR** Identify the gene-tissue pair that shares a causal signal with disease and estimate the corresponding cis-MR effect at a multi-signal locus. | 3 | 100.0% | Multi-signal conditional decomposition; ancestry-matched LD choice; colocalization; conditional cis-MR | Gene; tissue; colocated SNP; cis-MR effect | *gwas_sumstats*; eQTL panels (GENEA/GENEB by tissue); *study_metadata*; *gene_info*; ancestry-specific LD matrices |
| **Cross-platform pQTL colocalization and conditional cis-MR** Disentangle affinity artifacts, assay-specific binding effects, and shared signals across three assays and a disease GWAS. | 7 | 0.0% | Multi-signal conditional decomposition; cross-platform affinity contrast; colocalization (ABF); representative variant selection; conditional cis-MR | Assay-specific variant; follow-up variant; log-OR per SD protein | *variant_info* (metadata); *trait_metadata* (assays); *ld_matrix*; *ld_variant_order*; *somascan*; *olink*; *massspec*; *disease_gwas* |
| **CRISPR screen fitness inference with guide confounding** Infer gene-level fitness effects from a pooled screen with seed toxicity, copy-number bias, nonlinear GC-content effects, censoring, and inactive guides. | 6 | 0.0% | Staged technical-bias regression; inactive-guide Bayes-factor detection; left-censored Tobit model; guide-level QC | Top depleted gene; second gene; direct log<sub>2</sub> depletion | *guide_counts* (per-guide counts); *guide_features* (GC, seed, CN); *gene_panel*; *screen_metadata* |
| **Pharmacogenomic time-to-event MSM** Estimate genotype-specific treatment hazard ratios from EHR data with time-varying treatment-confounder feedback. | 6 | 75.0% | Treatment-confounder feedback; person-interval expansion; stabilized IPTW; pooled logistic MSM; wash-in lag specification | Responder genotype; HR noncarriers; HR carriers | *patients* (genotypes, outcomes, covariates); *labs* (longitudinal biomarker); *data_dictionary* |
| **Population, quantitative, and microbial genetics** |  |  |  |  |  |
| **ARG-based recombination hotspot detection** Localize a recombination hotspot and estimate its intensity from local ARG segments despite formatting inconsistencies, unit mismatches, and low-support trees. | 4 | 94.3% | Posterior filtering; unit normalization; adjacent-tree collapse; breakpoint KDE; branch-length weighting | Hotspot center (bp); hotspot multiplier | *local_trees* (intervals, tree summaries); *region_info* (chromosome length) |
| **Two-pulse admixture timing** Estimate admixture times and admixture proportion from local-ancestry tract lengths under censoring and genetic map errors. | 4 | 12.5% | Mixture-of-exponentials fitting; right-censoring correction; phase-switch fragment merging; genetic map unit QC; left-truncation | Recent time (generations); ancient time (generations); recent pulse weight | *segments* (tract calls, posteriors); *genetic_map* (2 swapped chroms); *chrom_lengths* |
| *Population, quantitative, and microbial genetics* |  |  |  |  |  |
| **Direct and genetic nurture PGS effects** Estimate direct and genetic nurture PGS effects from transmitted and non-transmitted alleles under assortative mating with incomplete trios. | 6 | 0.0% | Family PGS decomposition; transmitted / non-transmitted separation; assortative mating correction; duo-family handling; missing-data encoding | Direct PGS effect ($\beta_{\text{direct}}$) | *trios.npy* (child, mother, father genotypes); *weights* (PGS weights); *phenotypes*; *family_meta* (duo flags, sibship); *marker_qc* |
| **Cortisol SNP heritability** Estimate cortisol SNP heritability after recovering noisy collection-site labels and adjusting for multi-ancestry population structure. | 4 | 88.9% | Phenotype transform; center recovery from metadata; recomputed PCs; variance-component heritability estimation; center-linked ancestry interaction control | SNP heritability ($h^2$) | *genotypes* (dosages); *samples*; *variants*; *phenotype*; *covariates* (PC1, demographics, noisy center metadata) |
| **Metagenomic differential abundance and strain deconvolution** Identify differentially abundant microbial species and estimate strain-mixture proportions under compositional bias and batch effects. | 6 | 16.7% | Spike-in normalization; mock-community calibration; species selection; strain-panel orientation; batch-specific allele-flip correction | Case-associated species; $\log_2$ fold-change; strain-A fraction (cases); strain-A fraction (controls) | *species_counts* (sample-by-species); *sample_metadata*; *sample_qc*; *ref_counts*; *ref_qc*; strain panels |
| *Clinical screening and liquid biopsy* |  |  |  |  |  |
| **NIPT fetal-fraction estimation and mosaic trisomy detection** Estimate fetal fraction and detect mosaic trisomy in cfDNA with allele-biased SNPs, GC bias, and a maternal CNV. | 4 | 0.0% | Allelic-bias SNP filtering; target-chromosome exclusion; GC correction; maternal CNV detection | Karyotype; FF; coverage shift ($\delta$); mosaic fraction | *informative_snps* (allele counts); *test_counts* (sample bins); *panel_counts* (controls); *bin_metadata*; *analysis_manifest* |
| **Tumor-versus-CH cfDNA deconvolution** Separate tumor-derived variants from clonal hematopoiesis-derived variants in plasma cfDNA and estimate tumor fraction. | 7 | 50.0% | Stratum-aware PON calling; simplex/duplex artifact QC; fragment-based tumor-vs-CH classification; tumor-fraction estimation | Tumor-locus count; CH-locus count; tumor fraction | *control_counts* (control by locus); *case_counts* (sample by locus); *molecule_profiles* (fragment bins); *locus_catalog* |
| **cfDNA methylation deconvolution** Estimate tumor fraction and tumor-specific methylation profiles from plasma cfDNA using matched leukocytes and a biased reference atlas. | 7 | 0.0% | Control-locus QC; separate nonconversion estimation; patient-specific background choice; marker restriction; reference calibration; back-calculation | Nonconversion rate; tumor fraction; tumor methylation at region R17 | *plasma_counts* (cfDNA); *leukocyte_counts* (matched normal); *reference_regions* (atlas) |
| *Cancer and immunogenomics* |  |  |  |  |  |
| **APOBEC mutational signature attribution** Estimate the effect of genotype on APOBEC-associated mutational signatures after removing FFPE artifacts and kataegic clusters. | 4 | 63.0% | Mutation-level QC; kataegis detection; signature attribution; confounder-adjusted regression | APOBEC logit effect | *mutations* (per-SNV calls); *sample_metadata*; *signature_profiles* |
| **Presentation-competent clonal neoantigen burden** Identify the tumor with the highest presentation-competent clonal neoantigen burden under HLA loss, germline leakage, and subclonality. | 5 | 0.0% | FFPE artifact detection; LOH-germline filtering; integrated DNA/RNA HLA competence; expression gating; clonality estimation | Highest-burden tumor; clonal burden; total neoantigens | *sample_manifest* (tumor purity); *hla_status*; *somatic_calls*; *expression_by_variant*; *binding_predictions* |
| *Cancer and immunogenomics* |  |  |  |  |  |
| **HRD genomic scar scoring** Compute HRD genomic scar scores from allele-specific copy number despite WGD misclassification and segmentation noise. | 6 | 5.7% | Copy-number-scale inference; telomere/centromere masking; segment merging; ploidy-adjusted LST; scar scoring | GIS per sample; HRD status | *segments* (allele-specific CN); *snps* (BAF); *sample_metadata*; *chrom_lengths*; *masks* |
| *Functional, deconvolution, and spatial genomics* |  |  |  |  |  |
| **Perturb-seq effects of STAT1 knockdown** Estimate the effect of STAT1 knockdown on IFN response and STAT1 expression under ambient guide contamination and perturbation escape. | 4 | 0.0% | Ambient-aware singlet calling; escape filtering; batch-phase standardization; gene-set scoring | IFN-response effect; STAT1 expression effect | *guide_counts* (per-cell guides); *empty_guide_counts* (ambient); *expression_counts*; *cell_meta*; *guide_map*; *gene_sets* |
| **Bulk RNA-seq deconvolution** Estimate the genotype effect on dendritic cell fraction from bulk RNA-seq under reference-panel mismatch, site confounding, and outlier samples. | 5 | 9.8% | Calibrator-based scaling; high-variance gene filtering; eQTL-marker exclusion; outlier detection; site-adjusted regression | Target cell type; logit effect | *bulk_expression*; *scrna_reference* (single-cell reference); *sample_metadata* (site, genotype); *calibration_cell_fractions* |
| **Spatial tumor cis-eQTL mapping** Identify the malignant-cell-autonomous cis-eQTL from spatial transcriptomics in the presence of spot swapping, neighborhood effects, and CN artifacts. | 4 | 2.3% | Spot-swapping correction; deconvolution; tumor-dominant spot selection; CN adjustment; candidate classification | Target gene; direct effect; context gene; artifact gene | *spot_counts*; *spot_meta*; *donor_variant*; *reference_profiles*; *spot_auxiliary*; *candidate_genes* |
| *Other applications* |  |  |  |  |  |
| **Variant penetrance in panel sequencing under verification bias** Estimate the effect of a variant on confirmed diagnosis from clinical panel sequencing under kit-specific dropout and nonrandom case review. | 5 | 0.0% | Kit-specific calibration (binomial mixture); IPW for verification bias; collider avoidance; logistic regression; risk prediction | Carrier log-OR ($\beta_{\text{carrier}}$); predicted risk for female carrier at $age_z = 0$, $pc1 = 0$ | *people* (roster, partial diagnoses, covariates); *counts* (focal-site counts); *controls* (kit-specific wells) |
| **Low-template SNP mixture kinship inference** Estimate POI mixture weight and sibling-versus-unrelated LR from a low-template SNP mixture with degraded replicate profiles. | 4 | 0.0% | All-state $\tau$ calibration; blank-locus QC; degraded-replicate detection; per-locus mixture likelihood; kinship LR | POI mixture weight; $\log_{10}$ LR (sibling vs. unrelated) | *panel_manifest* (AFs, efficiency); *poi_genotypes*; *blank_controls*; *control_replicates*; *mixture_raw*; *poi_support_calls* (deconvolution export) |
| **Microexon PSI estimation** Estimate PSI for a 16bp microexon under read-length bias, cryptic splice donors, and condition–batch imbalance. | 6 | 9.6% | Sample QC; unannotated junction discovery; cryptic donor detection; read-length bias modeling; calibration extrapolation; batch correction | PSI (control); PSI (treated); ΔPSI | *sample_metadata* (condition, batch, read length); *event_catalog*; *junction_meta*; *junction_counts*; *annotation.gtf.gz* |

**Supplementary Table 2:** Values underlying Figure 4. Overall pass rate is the unweighted mean of per-problem pass rates across the 103 benchmark problems. The 95% confidence intervals match **Figure 4A**. The regime columns match **Figure 4B**. Avg. tokens reports mean tokens used, computed as the number of tokens in the model’s full chain-of-thought trace and final response, excluding tool calls, rounded to the nearest 0.1k. Token counts are not directly comparable across the two model groups: the non-GPT models were accessed via OpenRouter and their token totals reflect OpenRouter accounting, while the GPT variants were accessed via internal tooling and their totals reflect the internal accounting. Token totals are omitted for the Pro runs. Replicate summaries report the mean, minimum, and maximum numbers of valid runs contributing to each model-problem pass rate. GPT-family rows report reasoning effort, with GPT-5 shown from none through high and later mainline GPT models shown from none through xhigh.

| Model setting | Mean | 95% CI | 0% | >0–10% | >10–50% | ≥50% | Avg. tokens | Mean reps | Min | Max |
| --- | --- | --- | --- | --- | --- | --- | --- | --- | --- | --- |
| MiMo V2 Pro | 1.6% | [0.3, 3.8] | 89.3% | 9.7% | 0.0% | 1.0% | 20.5k | 20.0 | 20 | 20 |
| Kimi K2.5 | 1.8% | [0.6, 3.8] | 84.5% | 13.6% | 1.0% | 1.0% | 35.5k | 20.0 | 20 | 20 |
| Grok 4.20 (reasoning enabled) | 2.1% | [0.6, 4.3] | 87.4% | 7.8% | 3.9% | 1.0% | 11.6k | 20.0 | 20 | 20 |
| Qwen 3.6 Plus | 2.7% | [0.9, 5.3] | 81.6% | 14.6% | 1.9% | 1.9% | 57.6k | 20.0 | 20 | 20 |
| MiMo V2.5 Pro | 3.0% | [1.3, 5.4] | 75.7% | 16.5% | 6.8% | 1.0% | 38.7k | 20.0 | 19 | 20 |
| GLM 5.1 | 4.2% | [2.1, 6.8] | 72.8% | 17.5% | 7.8% | 1.9% | 95.5k | 20.0 | 20 | 20 |
| Kimi K2.6 | 7.4% | [4.1, 11.4] | 65.0% | 21.4% | 8.7% | 4.9% | 74.8k | 20.0 | 20 | 20 |
| Gemini 3.1 Pro (high) | 11.2% | [7.2, 15.7] | 55.3% | 19.4% | 16.5% | 8.7% | 23.5k | 40.0 | 40 | 40 |
| GPT-5 (none) | 1.9% | [0.5, 4.1] | 87.4% | 9.7% | 1.9% | 1.0% | 2.8k | 25.0 | 25 | 25 |
| GPT-5 (low) | 1.8% | [0.6, 3.4] | 79.6% | 17.5% | 1.9% | 1.0% | 5.6k | 36.6 | 24 | 53 |
| GPT-5 (medium) | 2.5% | [1.1, 4.5] | 74.8% | 19.4% | 4.9% | 1.0% | 10.0k | 51.0 | 37 | 59 |
| GPT-5 (high) | 3.5% | [1.6, 6.0] | 73.8% | 17.5% | 6.8% | 1.9% | 15.9k | 25.0 | 24 | 25 |
| GPT-5.2 (none) | 1.7% | [0.3, 4.1] | 90.3% | 5.8% | 2.9% | 1.0% | 1.0k | 25.0 | 24 | 25 |
| GPT-5.2 (low) | 2.3% | [0.6, 4.7] | 85.4% | 10.7% | 2.9% | 1.0% | 4.9k | 25.0 | 24 | 25 |
| GPT-5.2 (medium) | 4.0% | [1.7, 7.0] | 78.6% | 11.7% | 7.8% | 1.9% | 12.0k | 25.0 | 24 | 25 |
| GPT-5.2 (high) | 5.8% | [3.1, 9.1] | 66.0% | 20.4% | 11.7% | 1.9% | 15.5k | 39.7 | 32 | 40 |
| GPT-5.2 (xhigh) | 9.4% | [5.8, 13.6] | 55.3% | 23.3% | 14.6% | 6.8% | 37.6k | 20.4 | 14 | 25 |
| GPT-5.4 (none) | 2.0% | [0.5, 4.4] | 88.3% | 6.8% | 3.9% | 1.0% | 1.6k | 25.0 | 24 | 25 |
| GPT-5.4 (low) | 4.3% | [2.3, 6.6] | 70.9% | 15.5% | 12.6% | 1.0% | 9.8k | 25.0 | 24 | 25 |
| GPT-5.4 (medium) | 8.9% | [6.1, 12.2] | 47.6% | 28.2% | 21.4% | 2.9% | 19.4k | 49.9 | 48 | 50 |
| GPT-5.4 (high) | 16.0% | [11.1, 21.6] | 50.5% | 17.5% | 19.4% | 12.6% | 21.2k | 25.0 | 24 | 25 |
| GPT-5.4 (xhigh) | 19.0% | [13.3, 25.0] | 49.5% | 14.6% | 20.4% | 15.5% | 36.4k | 24.9 | 24 | 25 |
| GPT-5.5 (none) | 1.9% | [0.5, 4.2] | 90.3% | 4.9% | 3.9% | 1.0% | 0.6k | 25.0 | 24 | 25 |
| GPT-5.5 (low) | 3.2% | [1.1, 6.0] | 85.4% | 7.8% | 4.9% | 1.9% | 5.3k | 24.9 | 24 | 25 |
| GPT-5.5 (medium) | 9.2% | [5.7, 13.2] | 59.2% | 18.4% | 15.5% | 6.8% | 13.7k | 25.0 | 24 | 25 |
| GPT-5.5 (high) | 22.2% | [16.1, 28.6] | 40.8% | 20.4% | 17.5% | 21.4% | 17.7k | 48.1 | 29 | 59 |
| GPT-5.5 (xhigh) | 25.0% | [18.5, 31.9] | 41.7% | 15.5% | 18.4% | 24.3% | 24.8k | 54.3 | 39 | 60 |
| GPT-5 Pro | 4.0% | [1.7, 7.0] | 68.0% | 26.2% | 2.9% | 2.9% | – | 39.2 | 25 | 40 |
| GPT-5.2 Pro | 10.8% | [6.4, 15.6] | 60.2% | 19.4% | 11.7% | 8.7% | – | 31.4 | 16 | 42 |
| GPT-5.4 Pro | 25.6% | [18.6, 32.8] | 51.5% | 7.8% | 14.6% | 26.2% | – | 20.0 | 20 | 20 |
| GPT-5.5 Pro | 33.2% | [25.1, 41.5] | 49.5% | 6.8% | 10.7% | 33.0% | – | 19.6 | 16 | 20 |

**Listing 1:**
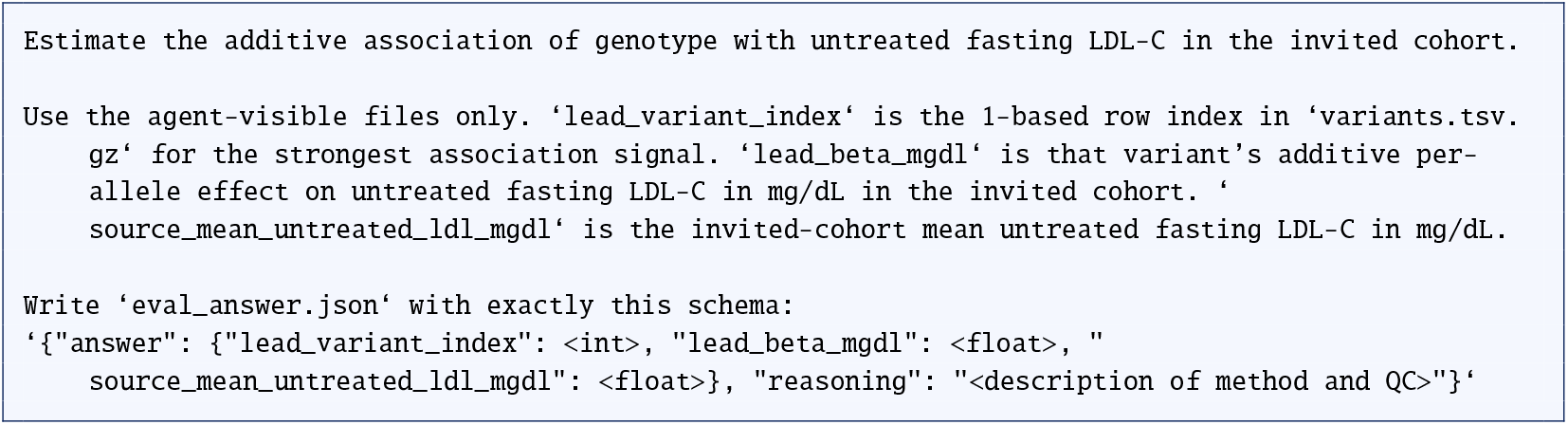
Evaluation prompt for the LDL GWAS follow-up case study.

## Appendix: LDL GWAS Case Study

This appendix illustrates the data generation process, correct approach, and ablation pipeline for one GeneBench problem. The task is a GWAS follow-up problem in which the goal is to identify a lead LDL variant and report its effect on untreated fasting LDL-C in the invited cohort. The problem is nontrivial because the primary, visible fasting-lab phenotype is both treatment-distorted and observed only in a selected follow-up subset, so a naive GWAS on the available lab values targets the wrong quantity in the wrong population.

Low-density lipoprotein cholesterol (LDL-C) is one of the most extensively studied quantitative traits in cardiovascular genetics. Large GWAS of blood lipids have identified many loci associated with LDL-C and related traits. ^37^ These associations are frequently used to motivate downstream biological and therapeutic investigation. ^38–40^ LDL-C is therefore a natural case study for benchmarked scientific analysis: the phenotype is clinically important, the association model is standard, and the downstream interpretation is decision-relevant.

At the same time, LDL-C analysis in real cohorts is often complicated by treatment and ascertainment. For instance, lipid-lowering therapy changes the measured phenotype, and treatment response is heterogeneous. ^44,46^ Refill histories can be more informative than self-report alone. ^45^ Selective participation can distort downstream association estimates. ^49,50^ The practical problem represented here reflects a common class of analyses in which the target estimand cannot be directly observed from the observed phenotype and the observed analytic subset and thus must be inferred through a series of corrective steps.

The problem retains the design principles of GeneBench that matter analytically: a minimal prompt, a recoverable target, simulated data, multi-stage inference with multiple decision points, threshold-robust QC, and ablation studies.

We first introduce the formal estimand and agent-visible files, the data-generating process, and then the three decision points required for valid estimation: reconstruction of untreated LDL-C, reweighting of attendees back to the invited cohort, and variant-level QC before the final scan. We close with the correct result and an ablation table showing how representative incorrect analyses fail.

### Problem Background and Estimand

The prompt provided to the agent for this problem is shown in Listing 1.

#### Agent-visible files

The agent-visible files are `cohort.tsv.gz, audit.tsv.gz`, and `variants.tsv.gz`. Together they define three linked views of the problem. The full cohort table covers all invited subjects and includes covariates, capillary LDL-C measured at invitation, treatment proxies, and fasting-lab LDL-C only for the subset who later return for a standardized fasting follow-up visit. The audit table can be joined back to the cohort by `sample_id`; it is then a random subset of those fasting-follow-up attendees for whom a historical untreated LDL-C measurement is also available.

The task is to identify which variant has the strongest additive association with *untreated* fasting LDL-C in the *invited cohort*. The cohort table contains 520 invited subjects with demographics, two ancestry principal components, travel distance, invitation wave, capillary LDL-C, fasting-lab LDL-C for attendees only, treatment proxies, attendance metadata, and genotype dosages at 60 variants. The cleaner fasting-lab phenotype is therefore not observed at invitation and is available only for the subset who later return for the standardized fasting follow-up visit. The audit table contains a random 100-subject validation sample of those fasting-lab attendees, each with a historical untreated LDL-C measurement. The variant manifest maps the genotype columns `v01`–`v60` to the 1-based variant indices used for reporting. These three files also map directly to the three analytic decision points: `audit.tsv.gz` supports calibration on the audited subset, `cohort.tsv.gz` supports attendance modeling on all invitees, and `cohort.tsv.gz` together with `variants.tsv.gz` support variant QC and final reporting.

#### Notation

Before turning to the estimand, it helps to fix the main objects once. Subjects are indexed by *i* and variants by *j*. We distinguish three nested subject sets: the invited cohort ℐ= {1, …, *N*}, the fasting-lab attendee subset 𝒜 = *i* : *A*_*i*_ = 1, and the audited attendee subset 𝒟 ⊆ 𝒜 for which historical untreated LDL-C is observed in `audit.tsv.gz`. At the phenotype level, *C*_*i*_ is capillary LDL-C measured at invitation, *L*_*i*_ is observed fasting-lab LDL-C among attendees, and *U*_*i*_ is the untreated fasting LDL-C target, observed only for *i* ∈ 𝒟 and latent elsewhere. The key treatment and selection variables are *R*_*i*_ (refill-based treatment proxy), 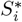 (self-reported statin use), and *A*_*i*_ (attendance at fasting follow-up). Later we write 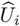 for the calibrated untreated-LDL prediction, *w*_*i*_ for the stabilized inverse-probability attendance weight, and 𝒥 _QC_ for the QC-passing variant set.

##### Target estimand

The scientific target motivating the problem is the invited-cohort additive association between genotype and untreated fasting LDL-C, together with the corresponding invited-cohort mean. Writing *U*_*i*_ for the target phenotype, *G*_*ij*_ for the genotype dosage, and *X*_*i*_ for the adjustment set,

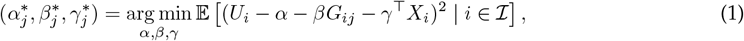

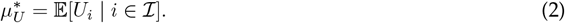

Since hidden untreated phenotypes and latent DGP parameters are not exactly recoverable through the agent-visible files, GeneBench grades the *recoverable realized-data target induced by the intended analysis path on the agent-visible files*. The graded path introduces four derived objects: 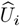, the audit-based prediction of untreated LDL-C for attendees; *w*_*i*_, the stabilized inverse-probability attendance weights that reweight fasting-lab attendees back to the invited cohort; 𝒥_QC_, the QC-passing variant set; and *p*_*j*_, the final weighted-association *p*-value for variant *j*. The graded lead variant is

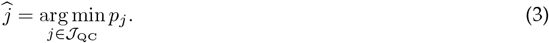

For each *j* ∈ 𝒥_QC_, the graded effect estimate is the genotype coefficient from

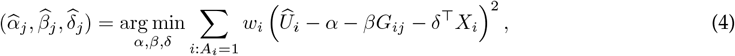

and the graded mean is

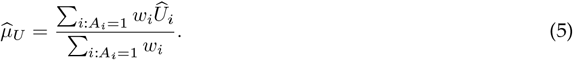

**Appendix Table 1:** Notation used in the LDL case study. Observed quantities map directly to agent-visible columns; derived or latent quantities are computed from those files or introduced during the analysis; standard notation and transforms are listed for reference.

| Symbol | Meaning | Agent-visible variable(s) |
| --- | --- | --- |
| *Observed quantities* |  |  |
| $i \in \mathcal{I} = \{1, \dots, N\}$ | Invitee index in the invited cohort, with $N = 520$. | rows |
| $j = 1, \dots, 60$ | Variants in `variants.tsv.gz`. | `variant_index`; v01–v60 |
| $G_{ij} \in \{0, 1, 2, \text{NA}\}$ | Additive genotype dosage for invitee $i$ at variant $j$; NA denotes a missing genotype call. | v01–v60 |
| $U_i$ | Untreated fasting LDL-C; observed only for $i \in \mathcal{D}$ and latent otherwise. | `baseline_ldl_mgdl` for audited attendees only |
| $L_i$ | Observed fasting-lab LDL-C; observed only when $A_i = 1$. | `lab_ldl_mgdl` |
| $C_i$ | Observed capillary LDL-C. | `capillary_ldl_mgdl` |
| $A_i$ | Indicator for attendance at the fasting-lab visit. | `attended_fasting_lab` |
| $S_i^*$ | Self-reported statin use. | `self_report_statin` |
| $R_i$ | Refill-based treatment proxy. | `refill_proxy` |
| $\text{dist}_i \in \mathbb{R}_+$ | Travel distance in km. | `dist_km` |
| *Derived or latent quantities* |  |  |
| $S_i$ | True statin exposure. | Not directly observed |
| $\text{wave}_i \in \{0, 1\}$ | Zero-based invitation wave, defined as the agent-visible `invite_wave` minus 1; thus $\text{wave}_i = 1$ corresponds to `invite_wave=2` in `cohort.tsv.gz`. | `invite_wave` |
| $\mathcal{A} = \{i : A_i = 1\}$ | Fasting-lab attendee subset. | `attended_fasting_lab` |
| $\mathcal{D} \subseteq \mathcal{A}$ | Audited attendee subset with observed $U_i$. | `sample_id` |
| $X_i$ | Association-model covariates: ($\text{age}_i, \text{sex}_i, \text{BMI}_i, \text{PC1}_i, \text{PC2}_i$). | `age`, `sex`, `bmi`, `pc1`, `pc2` |
| $Z_i$ | Attendance-model covariates: ($C_i, \text{age}_i, \text{BMI}_i, \text{sex}_i, \text{dist}_i, \text{wave}_i, \text{PC1}_i, S_i^*$). | `capillary_ldl_mgdl`, `age`, `bmi`, `sex`, `dist_km`, `invite_wave`, `pc1`, `self_report_statin` |
| $\hat{U}_i$ | Audit-calibrated prediction of untreated fasting LDL-C for attendee $i$. | `lab_ldl_mgdl`, `refill_proxy`, `self_report_statin`, `capillary_ldl_mgdl`, `age`, `sex`, `bmi`, `baseline_ldl_mgdl` |
| $\pi_i$ | Attendance probability $\Pr(A_i = 1 \mid Z_i)$. | `attended_fasting_lab`, `capillary_ldl_mgdl`, `age`, `bmi`, `sex`, `dist_km`, `invite_wave`, `pc1`, `self_report_statin` |
| $w_i$ | Stabilized inverse-probability attendance weight $\bar{A}/\hat{\pi}_i$. | `attended_fasting_lab`; $\hat{\pi}_i$ |
| $\mathcal{J}_{QC}$ | Variant set passing attendee-subset call-rate and Hardy–Weinberg filters. | v01–v60 |
| $p_j$ | Final weighted-association $p$-value for variant $j$. | $\hat{U}_i, G_{ij}, X_i$ |
| *Standard notation and transforms* |  |  |
| $\mathbf{1}(\cdot)$ | Indicator function. | – |
| $z(C_i) = (C_i - \bar{C})/s_C$ | Standardized capillary LDL-C in the realized invited cohort, where $\bar{C}$ and $s_C$ are the empirical mean and standard deviation of $C$. | `capillary_ldl_mgdl` |
| $Q_q(C)$ | Empirical $q$-quantile of $C$. | `capillary_ldl_mgdl` |
| $\min\{\max\{x, a\}, b\}$ | Truncation of $x$ to the interval $[a, b]$. | – |
| $\hat{\cdot}$ | Estimated quantity computed from the agent-visible files. | – |

These quantities are defined in detail below; the values against which outputs are graded (i.e. the ground truth values) are variant 42, 9.96 mg/dL, and 123.09 mg/dL, with grading tolerances of *±*0.40 mg/dL for the effect estimate and *±*1.00 mg/dL for the mean.

#### Data-Generating Process

The data-generating process was constructed so that the target remains recoverable from the observed data, while plausible partial analyses remain quantitatively wrong. Genotypes are additive dosages at 60 variants:

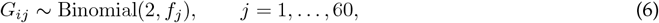

with *f*_*j*_ ∈ [0.08, 0.45] and mild PC1-linked frequency distortion for variants 6 and 13. Variant 42 is the sole causal locus:

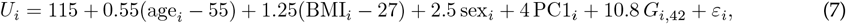

with *ε*_*i*_ ~ 𝒩 (0, 10^2^). This parameterization yields a single causal association of moderate size on the untreated phenotype scale. The 520-subject cohort stabilizes both the naive association scan and the induced attendance pattern. The 100-subject audit subset is sufficient to fit a multivariable calibration model without making untreated LDL-C effectively observed for the full cohort.

All 520 subjects also receive a noisier capillary proxy,

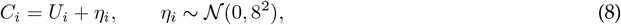

and treatment is assigned as a function of the latent untreated burden,

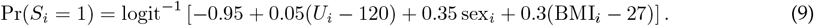

Medication intensity is summarized by a refill proxy. Let

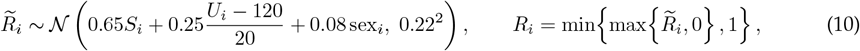

with refill-based medication summaries serving as a standard proxy for statin adherence in observational settings. ^45^ Self-report is deliberately imperfect:

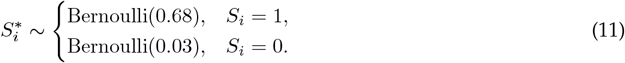

The observed fasting-lab phenotype is then

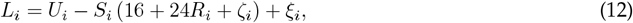

with *ζ*_*i*_ ~ 𝒩 (0, 4^2^) and *ξ*_*i*_ ~ 𝒩 (0, 6^2^). These parameters induce treatment effects that vary continuously with refill intensity rather than a single treated-versus-untreated offset. Under this construction, exclusions and flat offsets are mis-specified, whereas regression calibration remains the natural approach to the problem. ^41^ More general work on treatment-distorted quantitative traits also argues against naive exclusions or simple treated-versus-untreated adjustments, ^44^ and heterogeneous LDL response to statin therapy is well documented in pharmacogenetic studies. ^46^

Attendance at the fasting-lab visit is also non-random:

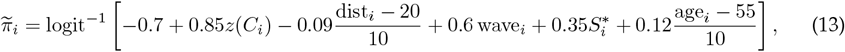

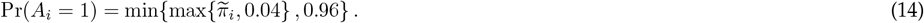

Attendance is therefore more likely among subjects with higher capillary LDL-C, older age, shorter travel distance, later invitation wave, and self-reported statin use. In the realized data, attendance rises from 15% in the lowest capillary-LDL quintile to 76% in the highest, and the lower clipping bound is active in the realized simulation. The resulting selection problem is substantial but identifiable. ^49,50^ Invited-cohort inference therefore requires inverse-probability weighting rather than ad hoc imputation of outcomes for non-attendees. ^42,43^

Finally, two technical QC failures are built directly into the genotype panel:

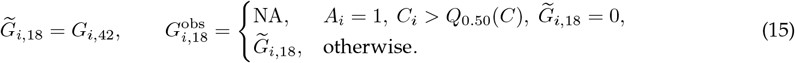

and

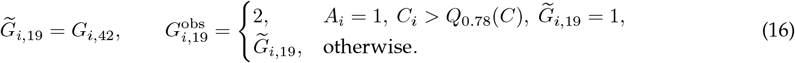

Variant 18 therefore behaves like a degraded proxy of the true signal with allele-specific dropout among higher-LDL attendees, whereas variant 19 behaves like a second degraded proxy with phenotype-linked heterozygote inflation. In the realized sample, variant 18 has attendee call rate 0.85 and variant 19 has HWE *p* = 5.1 × 10^−7^, so a standard QC pass now requires both the call-rate and Hardy–Weinberg filters. ^47,48^

#### Decision Point 1: Reconstruct Untreated LDL Before Scanning

Association analysis on observed fasting-lab LDL-C attenuates the target effect because the phenotype is measured on a treatment-distorted scale. In the realized data, the raw-lab scan still ranks variant 42 first, but the estimated per-allele effect is 4.56 mg/dL, well below the target value. **Appendix Figure 1** shows both the attenuation on the observed scale and the recovery after audit-based calibration.

**Appendix Figure 1:**
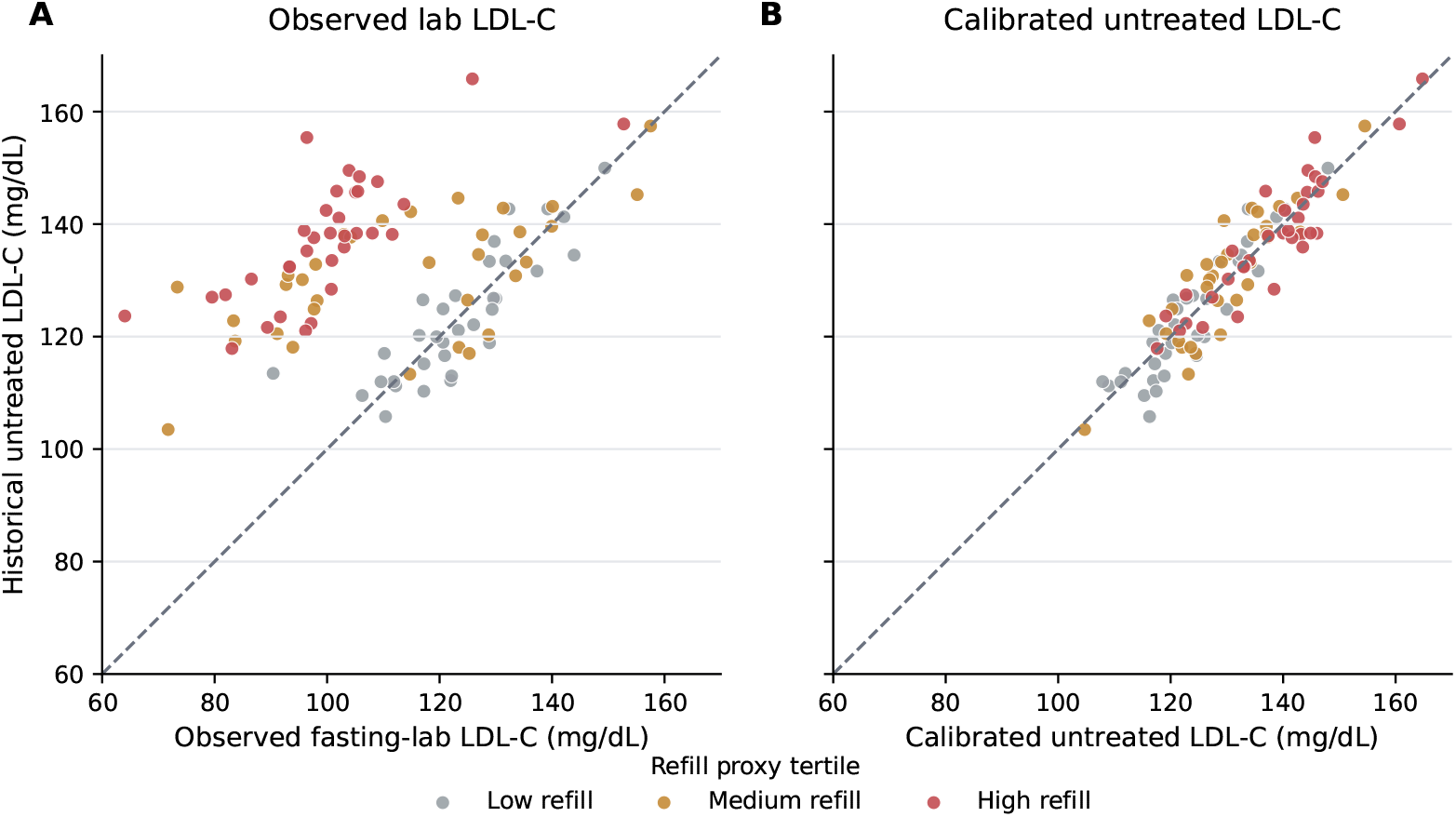
Decision point 1: treatment masking. **(A)** In the audit subset, observed fasting-lab LDL-C is compressed relative to historical untreated LDL-C, with the largest downward distortion in the highest refill tertile. **(B)** Audit-based regression calibration restores the untreated phenotype scale, bringing the calibrated values close to the identity line.

The audit subset provides the relevant diagnostic. Among audited subjects, using empirical tertiles of *R*_*i*_ (refill-based treatment proxy), the untreated-minus-lab gap is 0.12 mg/dL in the low-*R*_*i*_ tertile, 17.52 mg/dL in the middle-*R*_*i*_ tertile, and 37.31 mg/dL in the high-*R*_*i*_ tertile. The treatment effect is therefore heterogeneous and not well represented by a binary treated-versus-untreated adjustment.

##### Identifying the calibration predictor set

GeneBench problems are designed to be insensitive to nearby defensible analyst choices and sensitive only to missing scientifically necessary stages (Section 2); the calibration predictor set is a concrete illustration. The prompt does not specify which audit predictors to use; the agent must identify them from the data. Nested *R*^2^ on the 100-subject audit converges on a minimum-sufficient set {*L*_*i*_, *R*_*i*_, *C*_*i*_} : with demographics (age_*i*_, sex_*i*_, BMI_*i*_) always included, adding *L*_*i*_ alone gives *R*^2^ = 0.27, *C*_*i*_ alone gives *R*^2^ = 0.77, and further adding *R*_*i*_ to {*L*_*i*_, *C*_*i*_} raises *R*^2^ to 0.848. Each of the three contributes at least 0.04 in audit *R*^2^ and is individually significant at *p <* 0.001 in the full model, while 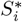 and each demographic covariate contribute less than 0.01 in audit *R*^2^ and are not individually significant at *p <* 0.05. All specifications that retain {*L*_*i*_, *R*_*i*_, *C*_*i*_} land inside the graded tolerance, while dropping any one of the three fails (**Appendix Table 2**). The visual diagnostic in **Appendix Figure 1A** reinforces this: subjects with similar *L*_*i*_ separate vertically by *R*_*i*_, so *L*_*i*_ alone does not recover *U*_*i*_, and the remaining spread within refill strata is explained by *C*_*i*_, which tracks untreated burden before treatment compression.

Untreated LDL-C is reconstructed by regression calibration, using the 100 audited attendees as a validation sample. ^41^ The working model is

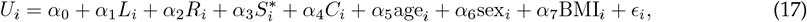

fit in the audit subset and then used to predict 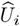 for all attendees. The full model achieves in-sample audit *R*^2^ = 0.85 and includes 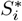 and demographics as conservative additions beyond the minimum-sufficient surrogate set {*L*_*i*_, *R*_*i*_, *C*_*i*_}. **Appendix Table 2** confirms this decomposition. Specifications that retain all of {*L*_*i*_, *R*_*i*_, *C*_*i*_} land inside the graded tolerance: dropping 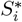 gives 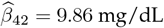 mg/dL, dropping demographics gives 10.27, and the minimal *L*_*i*_, *R*_*i*_, *C*_*i*_-only calibration gives 10.20. Specifications that drop any one of {*L*_*i*_, *R*_*i*_, *C*_*i*_} fail: removing *R*_*i*_ yields 9.28, removing *C*_*i*_ yields 8.15, using *L*_*i*_ plus demographics alone yields 1.66, and a flat 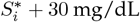 correction yields 7.77. In the realized data, calibration residuals also remain centered across the fitted range, so the agent-visible problem does not require a more elaborate nonlinear working model.

#### Decision Point 2: Reweight Attendees Back to the Invited Cohort

Phenotype reconstruction does not by itself restore the target population. The unweighted attendee mean of calibrated untreated LDL-C is 129.65 mg/dL, whereas the invited-cohort target is 123.09 mg/dL. The discrepancy reflects selection into the fasting-lab visit. **Appendix Figure 2** shows the resulting distortion and the population transport achieved by inverse-probability weighting. The figure uses *C*_*i*_ (capillary LDL-C at invitation) because it is observed for all invitees and is a principal driver of attendance, whereas *L*_*i*_ and *U*_*i*_ are not available for the full invited cohort. This is therefore a standard selection-on-observables problem: the outcome is available only for attendees, but attendance depends on invitation-time covariates observed for the full cohort. Under that structure, inverse-probability weighting is the natural transport estimator because it reweights observed attendee outcomes back to the invited-cohort covariate distribution without imputing untreated LDL-C for non-attendees. ^42,43^

**Appendix Figure 2:**
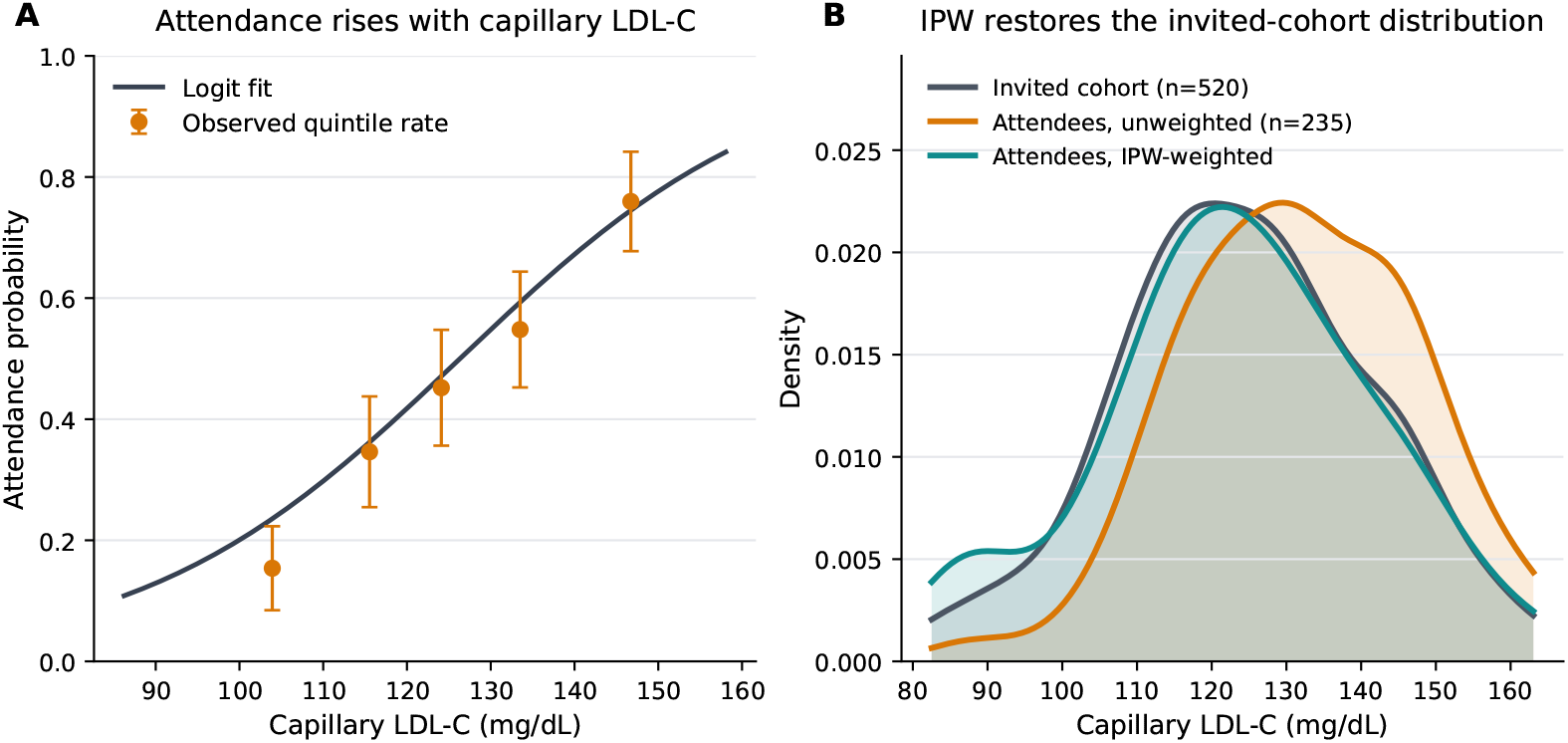
Decision point 2: selective follow-up. **(A)** Attendance probability rises sharply with capillary LDL-C, so the fasting-lab subset is enriched for higher-LDL invitees. **(B)** The unweighted attendee capillary-LDL distribution is shifted relative to the invited cohort, whereas inverse-probability weighting maps it back toward the invited-cohort distribution. Capillary LDL-C is shown because it is observed at invitation for the full cohort and enters the attendance model directly.

Analyses that stop after calibration therefore remain targeted to the wrong population. In the realized data, calibrated but unweighted analysis gives 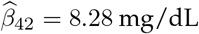 and overestimates the mean untreated LDL-C by 6.56 mg/dL. The attendance model must be fit on all 520 invitees, because attendance is defined for the full invited cohort. Untreated LDL-C is predicted only for attendees, where the observed lab phenotype and audit relationship make that reconstruction identifiable; the weights then transport the attendee analysis back to the invited cohort.

Stabilized inverse-probability weights are therefore estimated on all 520 invitees. ^42,43^ Let

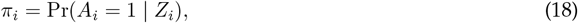

with logistic working model

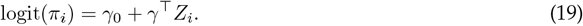

We use a logistic propensity model because attendance is binary and the benchmark’s selection mechanism is designed to be well approximated by a low-dimensional logistic model. The stabilized weights are

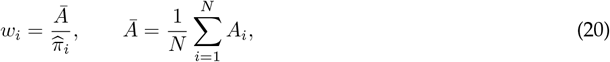

with 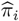 truncated to [0.05, 0.95]. The resulting stabilized weights are then truncated at the 1st and 99^th^ percentiles of the full invited-cohort weight distribution before restriction to attendees for the outcome analysis. In the realized sample, the attendee weights have median 0.78 and 99th percentile 5.05, with maximum 7.01. The weights are therefore large enough to shift the target population but not so extreme that the result is driven by trimming choice. Note that the percentile truncation is primarily a safeguard against extreme weights, but in this particular problem, lack of truncation does not materially change the result and leaves the graded outputs within tolerance. Outcome prediction is applied only to attendees. The audit subset is therefore used for phenotype reconstruction, not for the attendance model itself.

#### Decision Point 3: Perform Variant QC Before Ranking Associations

Variant-level QC remains necessary after phenotype reconstruction and reweighting. In this benchmark, variant 18 is a degraded proxy of the true signal with allele-specific dropout among higher-LDL attendees, and variant 19 is a second degraded proxy with phenotype-linked heterozygote miscoding. **Appendix Figure 3** shows that the realized instance now requires both QC filters before association ranking.

**Appendix Figure 3:**
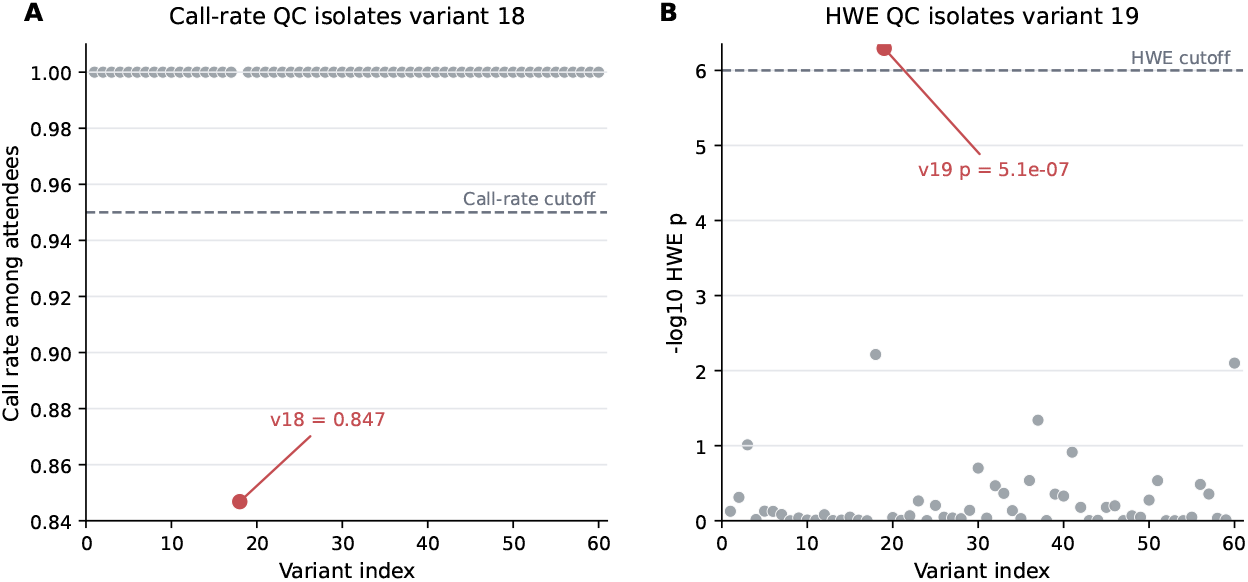
Decision point 3: variant QC. Variant 18 exhibits phenotype-linked missingness and fails call-rate QC, while variant 19 preserves call rate but fails Hardy–Weinberg equilibrium. Full QC requires both the call-rate and Hardy–Weinberg filters before the final association scan.

QC is implemented on the attendee subset used for association testing, with standard call-rate and Hardy– Weinberg thresholds,

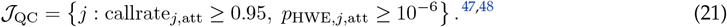

In the realized data, variant 18 has attendee call rate 0.85 and HWE *p* = 0.006, while variant 19 has attendee call rate 1.00 but HWE *p* = 5.1 × 10^−7^. Every other variant clears both filters. If QC is skipped, variant 18 rather than variant 42 becomes the lead signal, with 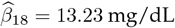 and *p* = 4.3 × 10^−25^. If the analyst filters only on call rate and skips HWE, variant 19 becomes the lead instead, with 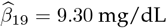 and *p* = 2.3 × 10^−22^.

#### Correct Result and Ablations

After phenotype reconstruction, reweighting, and attendee-subset QC, the final association scan fits a weighted additive regression on the attendees,

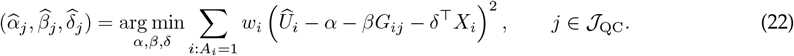

The lead variant is the QC-passing variant with the smallest corresponding *p*-value. The corresponding invited-cohort mean is

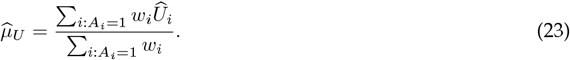

In the realized dataset, variant 42 is the lead signal with 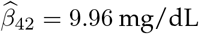, *p* = 2.7 × 10^−18^, and 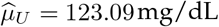.

**Appendix Table 2:** Ablation results for the intended analysis and representative wrong approaches. The top block lists calibration specifications that pass the graded tolerance; all retain the minimum-sufficient surrogate set {*L*_*i*_, *R*_*i*_, *C*_*i*_}. The remaining rows drop one of those surrogates or skip a decision point, and each fails on at least one graded quantity even when the lead variant is still ranked correctly. Values are rounded to two decimal places for presentation.

| Analysis | Pass? | Lead variant | Effect estimate (mg/dL) | Effect error (mg/dL) | Reported mean LDL-C (mg/dL) | Mean error (mg/dL) |
| --- | --- | --- | --- | --- | --- | --- |
| correct | Pass | 42 | 9.96 | 0.00 | 123.09 | 0.00 |
| calibrated_no_selfreport_weighted | Pass | 42 | 9.86 | -0.10 | 123.36 | 0.27 |
| calibrated_no_demographics_weighted | Pass | 42 | 10.27 | 0.32 | 123.36 | 0.27 |
| calibrated_minimal_LRC_weighted | Pass | 42 | 10.20 | 0.25 | 123.69 | 0.59 |
| raw_lab_unweighted | Fail | 42 | 4.56 | -5.39 | 110.48 | -12.61 |
| raw_lab_weighted | Fail | 42 | 6.70 | -3.26 | 109.18 | -13.92 |
| self_report_plus30_unweighted | Fail | 42 | 4.77 | -5.18 | 123.12 | 0.02 |
| self_report_plus30_weighted | Fail | 42 | 7.77 | -2.19 | 117.97 | -5.12 |
| calibrated_unweighted | Fail | 42 | 8.28 | -1.68 | 129.65 | 6.56 |
| calibrated_no_refill_weighted | Fail | 42 | 9.28 | -0.68 | 122.84 | -0.25 |
| calibrated_no_capillary_proxy_weighted | Fail | 42 | 8.15 | -1.81 | 124.93 | 1.84 |
| calibrated_lab_only_weighted | Fail | 42 | 1.66 | -8.29 | 127.85 | 4.76 |
| capillary_proxy_unweighted | Fail | 42 | 10.84 | 0.89 | 131.99 | 8.89 |
| capillary_proxy_weighted | Fail | 42 | 13.14 | 3.19 | 123.45 | 0.36 |
| calibrated_weighted_no_covariates | Fail | 42 | 10.63 | 0.67 | 123.09 | 0.00 |

**Appendix Table 2** summarizes the ablations. Naive lab analyses attenuate the effect estimate, and partially calibrated analyses recover some of the phenotype scale but remain biased. Unweighted calibrated analyses remain targeted to the attendee subset, and capillary-proxy analyses operate on the wrong phenotype scale. The benchmark therefore requires recovery of the correct estimand and of the sequence of upstream corrections required to estimate it.

